# Retrograde adenosine/A_2A_ receptor signaling facilitates excitatory synaptic transmission and seizures

**DOI:** 10.1101/2021.10.07.463512

**Authors:** Kaoutsar Nasrallah, Coralie Berthoux, Yuki Hashimotodani, Andrés E. Chávez, Michelle Gulfo, Rafael Luján, Pablo E. Castillo

**Affiliations:** Dominick P. Purpura Department of Neuroscience, Albert Einstein College of Medicine, Bronx, NY 10461, U.S.A; Department of Psychiatry & Behavioral Sciences, Albert Einstein College of Medicine, Bronx, NY 10461, U.S.A; Instituto de Investigación en Discapacidades Neurológicas (IDINE), Facultad de Medicina, Universidad Castilla-La Mancha, 02008 Albacete, Spain; Graduate School of Brain Science, Doshisha University, Kyoto, Japan; Centro Interdisciplinario de Neurociencia de Valparaíso, Facultad de Ciencias, Universidad de Valparaíso, Valparaíso 2340000, Chile; Department of Biological Sciences, Fordham University, Bronx, NY 10458, U.S.A

**Keywords:** Mossy cell, hippocampus, dentate gyrus, retrograde signaling, epilepsy, presynaptic, BDNF, TrkB, PKA, LTP

## Abstract

Retrograde signaling at the synapse is a fundamental way by which neurons communicate and neuronal circuit function is fine-tuned upon activity. While long-term changes in neurotransmitter release commonly rely on retrograde signaling, the mechanisms remain poorly understood. Here, we identified adenosine/A_2A_ receptor (A_2A_R) as a novel retrograde signaling pathway underlying presynaptic long-term potentiation (LTP) at a hippocampal excitatory circuit critically involved in memory and epilepsy. Transient burst activity of a single dentate granule cell induced LTP of mossy cell synaptic inputs, a BDNF/TrkB-dependent form of plasticity that facilitates seizures. Postsynaptic TrkB activation released adenosine from granule cells, uncovering a non-conventional BDNF/TrkB signaling mechanism. Moreover, presynaptic A_2A_Rs were necessary and sufficient for LTP. Lastly, seizure induction released adenosine in a TrkB-dependent manner, while removing A_2A_Rs or TrkB from the dentate gyrus had anti-convulsant effects. By mediating presynaptic LTP, adenosine/A_2A_R retrograde signaling may modulate dentate gyrus-dependent learning and promote epileptic activity.

**Highlights:**

- Postsynaptic firing induces presynaptic LTP at mossy cell to granule cell synapses
- Postsynaptic TrkB activation induces adenosine release from granule cells
- Presynaptic adenosine A_2A_ receptors are necessary and sufficient to induce LTP
- Adenosine/A_2A_R signaling within the dentate gyrus is pro-convulsant

**In Brief:** Nasrallah et al. report a novel retrograde signaling pathway at hippocampal synapses that involves postsynaptic TrkB-dependent release of adenosine and the activation of presynaptic A_2A_ receptors. This pathway mediates presynaptic long-term potentiation at a key hippocampal excitatory synapse and can also promote epileptic seizures.

## INTRODUCTION

Retrograde signaling at the synapse is a well-established mechanism whereby a neuron can strengthen or weaken the synaptic inputs it receives (Fitzsimonds and Poo, 1998; Regehr et al., 2009). Diverse types of messengers, including lipids, gases, peptides and conventional neurotransmitters, can be released from the postsynaptic neuron upon activity and activate presynaptic receptors or other molecular targets, establishing a retrograde signaling system that powerfully increases or decreases neurotransmitter release. Retrograde signaling has been implicated in short-term synaptic plasticity (Regehr *et al*., 2009), presynaptic long-term plasticity (Castillo, 2012; Monday et al., 2018) and also in presynaptic homeostatic plasticity (Davis and Muller, 2015), all of which rely on changes in neurotransmitter release. Despite the significance of retrograde signaling in fine-tuning neuronal circuit function, important knowledge gaps remain regarding the identity and release mechanism of the retrograde messenger, as well as the presynaptic targets and downstream molecular cascades implicated in regulating neurotransmitter release.

The dentate gyrus (DG), the major input area of the hippocampus, contains two main types of excitatory neurons: dentate granule cells (GCs) and hilar mossy cells (MCs). MCs and GCs form an associative circuit that is proposed to play a key role in DG-dependent cognitive functions (Lisman, 1999; Scharfman, 2016) and epilepsy (Botterill et al., 2019; Bui et al., 2018; Jinde et al., 2013; Nasrallah et al., 2022; Ratzliff et al., 2002; Scharfman and Myers, 2012). Repetitive stimulation of MC axons with physiologically relevant patterns of activity triggers robust presynaptic long-term potentiation at MC-GC synapses (MC-GC LTP) but not at MC to interneurons synapses (Hashimotodani et al., 2017). Recent evidence indicates that MC-GC LTP can be induced *in vivo* by enriched environment exposure (Berthoux et al., 2023) and experimental epileptic activity (Nasrallah *et al*., 2022). Uncontrolled strengthening of MC-GC transmission promotes seizures and likely contributes to the pro-epileptic role of MCs in early epilepsy stages (Botterill *et al*., 2019; Nasrallah *et al*., 2022). Thus, it is crucial to determine the precise mechanism underlying MC-GC LTP. This presynaptic form of plasticity is mechanistically unique. Its induction is NMDA receptor-independent but requires postsynaptic BDNF/TrkB signaling upstream of presynaptic cAMP/PKA signaling (Hashimotodani *et al*., 2017), strongly suggesting the involvement of a retrograde signaling system whose identity is unknown.

The following criteria must be satisfied to establish retrograde signaling as a mechanism of presynaptic long-term plasticity. First, the retrograde messenger must be synthesized and released from the postsynaptic compartment. Second, interfering with the synthesis and/or release of this messenger should prevent plasticity. Third, the target for the retrograde messenger must be present in the presynaptic bouton. Fourth, interfering with the presynaptic target should also prevent plasticity. Lastly, activation of the presynaptic target by the retrograde messenger or some analogous molecule should mimic long-term plasticity –although in some cases this activation alone may be insufficient to induce plasticity.

In this study, we sought to determine the retrograde signal involved in presynaptic LTP at MC-GC synapses. To our surprise, we found that most conventional retrograde messengers were not implicated in this form of plasticity. Using selective pharmacology, immunoelectron microscopy, and a conditional knockout strategy, we discovered that activation of presynaptic G_s_-coupled adenosine A_2A_ receptors (A_2A_Rs) is necessary and sufficient to induce MC-GC LTP. In addition, TrkB activation mobilized adenosine from GCs, uncovering a novel TrkB-dependent signaling mechanism, whereas interfering with GC adenosine release abolished LTP. Furthermore, adenosine was released *in vivo* during acutely induced epileptic seizures in a TrkB-dependent manner, while removing A_2A_Rs or TrkB from hippocampal excitatory neurons had anti-convulsant effects. Our findings not only establish adenosine/A_2A_R as a novel retrograde signaling system that mediates presynaptic plasticity, but also uncover a synapse-specific mechanism by which BDNF/TrkB and A_2A_Rs may promote epileptic activity.

## RESULTS

### Theta-burst firing of a single GC induces presynaptic LTP at MC-GC synapse

To identify the retrograde signal mediating MC-GC LTP, we first tested whether activation of a single GC, a manipulation previously used to characterize endocannabinoid retrograde signaling in long-term plasticity (Younts et al., 2013), could trigger MC-GC LTP. We found that GC theta-burst firing (TBF: 10 bursts at 5 Hz of 5 AP at 50 Hz, repeated 4 times every 5s, **Figure 1A**) induced LTP selectively at MC-GC synapses but not at neighboring medial perforant path (MPP)-GC synapses (**Figure 1B**). In addition, the group II mGluR agonist DCG-IV (1 µM), which selectively abolishes GC-MC (Lysetskiy et al., 2005) but not MC-GC synaptic transmission (Chiu and Castillo, 2008), did not affect the magnitude of TBF-induced LTP (**Figure S1A**), indicating that this plasticity does not rely on GC recruitment of MC firing. We then tested whether TBF-LTP shared the same mechanistic features as synaptically-induced MC-GC LTP. Like synaptically-induced MC-GC LTP (Hashimotodani *et al*., 2017), TBF-LTP is likely expressed presynaptically, as indicated by a significant reduction in both paired-pulse ratio (PPR) and coeCfficient of variation (CV) (**Figure 1C**) (see Methods). Moreover, TBF-LTP was abolished in the presence of the selective TrkB antagonist ANA-12 (15 µM) (**Figure S1B**), and in postsynaptic (**Figure 1D** and **Figure S1C**) but not presynaptic *TrkB* conditional knockout (cKO) mice (**Figure S1D**) (see Methods). TBF-LTP was also abolished in postsynaptic *Bdnf* cKO mice (**Figure S1E**), in GCs patch-loaded with botulinum toxin-B (Botox, 0.5 µM; **Figure S1F**), which blocks BDNF release (Shimojo et al., 2015), and in GCs loaded with the calcium chelator BAPTA (20 mM), which blocks postsynaptic calcium rise during TBF (**Figure S1G**). We discarded the possibility that alterations in GC membrane properties or basal transmitter release could underlie the lack of LTP in postsynaptic *Bdnf* cKO (Nasrallah et al., 2022) and postsynaptic *TrkB* cKO (**Figure S1H and S1I**). The PKA inhibitors H89 (10 μM, 40-to 60-min pre-incubation and bath applied) (**Figure S1J**) and membrane-permeable PKI_14–22_ myristoylated (1 μM, 40-to 60-min pre-incubation and bath applied)(**Figure 1E**) also abolished TBF-LTP, whereas loading the membrane-impermeable PKI_6-22_ (2.5 μM) in GCs did not affect this plasticity (**Figure 1E**). TBF-LTP was normally induced in the presence of the NMDAR antagonist D-APV (50 µM) (**Figure S1K**). Remarkably, while postsynaptic *TrkB* or *Bdnf* deletion abolished GC TBF-induced LTP (**Figure 1D,1F** and **Figure S1C, E**), it had no impact on the chemical LTP induced by transient activation of PKA with the adenylyl cyclase activator forskolin (Berthoux *et al*., 2023; Hashimotodani *et al*., 2017; Nasrallah *et al*., 2022), supporting the notion that PKA acts downstream of BDNF/TrkB. To remove *TrkB* from GC specifically, we took advantage of a lentivirus encoding Cre under the control of the C1ql2 promoter, which achieves sparse labeling of GCs only (**Figure S1C**) (Barthet et al., 2018; Monday et al., 2022). In all, these manipulations (summarized in **Figure 1F**) indicated that, like synaptically-induced MC-GC LTP, TBF-LTP requires postsynaptic BDNF/TrkB signaling and presynaptic PKA signaling and is NMDAR-independent (**Figure 1G**). Lastly, synaptically-induced MC-GC LTP and TBF-LTP occluded each other (**Figure S2**), indicating a common mechanism. Thus, GC TBF triggered a presynaptic form of LTP with identical properties to the synaptically induced MC-GC LTP (Hashimotodani *et al*., 2017), thereby establishing a simple, single-cell manipulation to investigate the identity of the retrograde messenger (**Figure 1G**).

**Figure 1:**
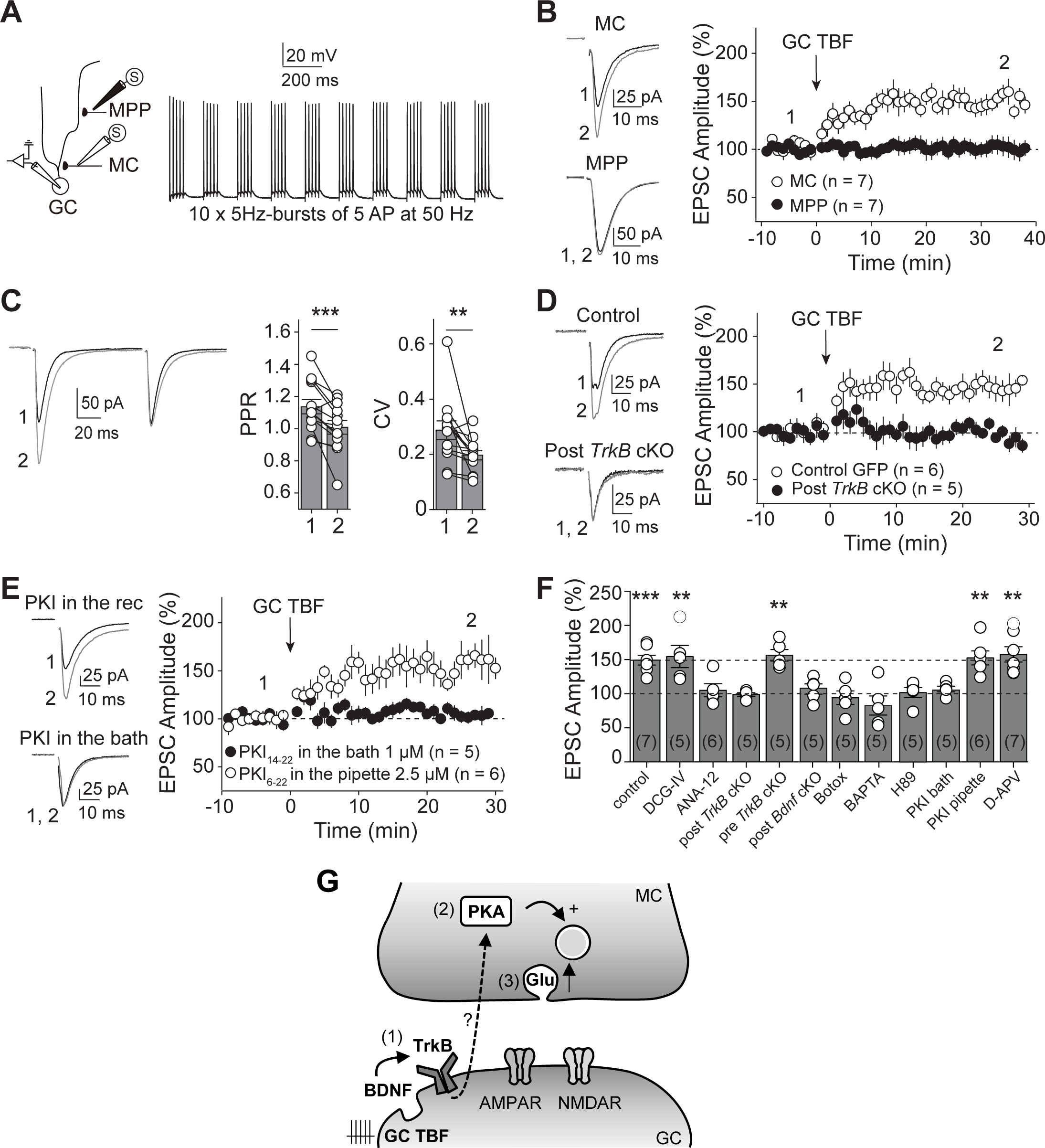
Theta-burst firing of a single GC induces presynaptic LTP at the MC-GC synapse. **(A)** *Left,* Diagram illustrating the recording configuration. MC and MPP EPSCs were recorded from the same GC and evoked with stimulation electrodes placed in the inner and middle molecular layer, respectively. *Right*, Current clamp recording showing GC theta-burst firing (GC TBF). LTP induction protocol (GC TBF) was composed of 10 bursts at 5 Hz of 5 action potentials at 50 Hz, repeated 4 times every 5 s. **(B)** *Left,* Representative traces before (1) and after (2) GC TBF delivery. *Right,* Time-course plot showing that GC TBF induced LTP at MC-GC but not at MPP-GC synapses. **(C)** GC TBF-induced LTP was associated with significant reduction in PPR and CV (n = 13 cells). ** p < 0.01, *** p < 0.001. **(D)** LTP was abolished when TrkB was conditionally knocked out from from GCs (Post *TrkB* cKO, *TrkB^fl/fl^* mice injected in the dorsal blade with AAV_5_.CaMKII.Cre.GFP). LTP was unaffected in control animals (Control, *TrkB_fl/fl_* mice injected in the dorsal blade with AAV_5_.CaMKII.eGFP). **(E)** LTP was normally induced when loading PKI_6-22_ (2.5 μM) in GCs via the recording pipette but completely blocked when the cell-permeable PKA inhibitor PKI_14-22_ myristoylated (1 μM) was bath applied. **(F)** Summary bar graph showing the magnitude of GC TBF-induced LTP in the presence of DGC-IV (1 μM), when TrkB was conditionally knocked out from MCs (Pre *TrkB* cKO), when loading the PKI_6-22_ (2.5 μM) in GCs, and in the presence of D-APV (50 µM). LTP was abolished in the presence of the TrkB antagonist ANA-12 (15 μM), when Botox (0.5 µM) was loaded postsynaptically, in postsynaptic BDNF and *TrkB* cKO mice, and during bath application of the PKA inhibitors H89 (10 μM) or myristoylated PKI_14-22_ (1 μM). Time-course summary plots are shown in Figure S1. ** p < 0.01, *** p < 0.001. **(G)** Scheme illustrating the emerging model for the mechanism underlying GC TBF-LTP. GC TBF triggers postsynaptic BDNF release and subsequent TrkB activation in GCs (1). Presynaptic PKA is then engaged downstream of postsynaptic BDNF/TrkB signaling (2), suggesting the requirement of a retrograde signal. Lastly, presynaptic PKA activation resulted in a long-lasting increase in glutamate release (3). Numbers in parentheses indicate the number of cells. Data are presented as mean ± SEM.

### A non-conventional retrograde signal likely mediates presynaptic LTP at MC-GC synapses

Previous work discarded endocannabinoids, glutamate, and GABA as retrograde signals mediating MC-GC LTP (Hashimotodani *et al*., 2017; Jensen et al., 2021), and our current finding showing that TBF-LTP was normally induced in presynaptic *TrkB* cKO mice (**Figure S1D**), also discards BDNF as retrograde messenger. We therefore tested the role of other conventional retrograde messengers, such as nitric oxide (NO) and lipid-derived messengers (Regehr *et al*., 2009). Application of the NO synthase inhibitor L-NAME (100 μM, 50-to 90-min pre-incubation and bath applied) did not impair TBF-LTP (**Figure S3A**), but as a positive control it significantly reduced LTP at CA3-CA1 synapses (**Figure S3B**), as previously reported (Williams et al., 1993). Blockade of lipid-derived messengers with a cocktail of inhibitors consisting of the fatty acid amide hydrolase (FAAH) and anandamide amidase inhibitor AACOCF3 (10 μM, 50-to 80-min pre-incubation and bath applied), the lipoxygenases inhibitor baicalein (3 μM, 50-to 80-min pre-incubation and bath applied), and the lipase inhibitor THL (tetrahydrolipstatin or orlistat, 4 μM in the recording pipette) did not impair TBF-LTP either (**Figure S3C**). As positive controls, we found that both AACOCF3 and baicalein significantly reduced long-term depression (LTD) at CA3-CA1 synapses (**Figure S3D**) (Normandin et al., 1996), and loading the lipase inhibitor THL (4 μM) in the recording pipette efficiently blocked CA1 inhibitory LTD (iLTD) (**Figure S3E**) (Chevaleyre and Castillo, 2003). Lastly, as was true for TBF-LTP, blocking NO signaling and lipid-derived messengers in interleaved control experiments did not alter synaptically-induced LTP either (**Figure S4**). Altogether, our results strongly suggest the involvement of a non-conventional retrograde messenger in presynaptic MC-GC LTP.

### Presynaptic A_2A_Rs are required for MC-GC LTP

Presynaptic PKA signaling is necessary and sufficient to induce LTP of MC-GC synaptic transmission (Hashimotodani *et al*., 2017), raising the possibility that the retrograde messenger induces LTP by activating a G_s_-coupled G protein-coupled receptor (GPCR) located on MC axon terminals. The G_s_-coupled adenosine A_2A_ receptor (A_2A_R) is a good candidate as it has been implicated in BDNF-mediated plasticity at other synapses (Diogenes et al., 2004; Fontinha et al., 2008; Sebastiao and Ribeiro, 2009b), and the endogenous ligand adenosine can be released from neurons upon activity (King et al., 2006; Latini and Pedata, 2001; Lovatt et al., 2012; Pons-Bennaceur et al., 2019; Wall and Dale, 2013). To test whether adenosine mediates MC-GC LTP by activating presynaptic A_2A_Rs, we first examined whether A_2A_R antagonism impaired this plasticity. Two different A_2A_R antagonists, SCH 58261 (100 nM) and ZM241385 (50 nM), abolished TBF-LTP (**Figure 2A**). Moreover, bath application of SCH 58261 did not change basal transmission (**Figure 2B**) but prevented the induction of LTP by BDNF puffs (8 nM, 2 puffs of 3 s) delivered in the inner molecular layer (IML) (**Figure 2C**) (Hashimotodani *et al*., 2017), strongly suggesting that A_2A_R activation is required downstream of the BDNF/TrkB cascade. It is unlikely that postsynaptic A_2A_Rs could be implicated given that including the GPCR inhibitor GDPβS (1 mM) in the recording pipette did not affect TBF-LTP, whereas as a positive control it abolished the change in holding current induced by the selective GABA_B_ receptor agonist baclofen (10 μM) (**Figure 2D**). Likewise, synaptically-induced MC-GC LTP was also abolished by SCH 58261 (**Figure S5A**), but not by including GDPβS in the recording pipette (**Figure S5B**). To demonstrate the role of presynaptic A_2A_Rs in MC-GC LTP, we also employed a conditional knockout strategy combined with optogenetics which allowed us to selectively activate A_2A_R-deficient MC axons expressing the fast opsin ChIEF (**Figure 2E**). We found that synaptic responses elicited by the activation of these axons did not undergo TBF-LTP (**Figure 2F**). In contrast, deleting *Adora2a* from MCs did not significantly affect basal neurotransmitter release, as indicated by the lack of PPR change (**Figure 2G**). Taken together, these findings indicate that presynaptic, and not postsynaptic, A_2A_R activation is necessary for LTP at MC-GC synapses.

**Figure 2:**
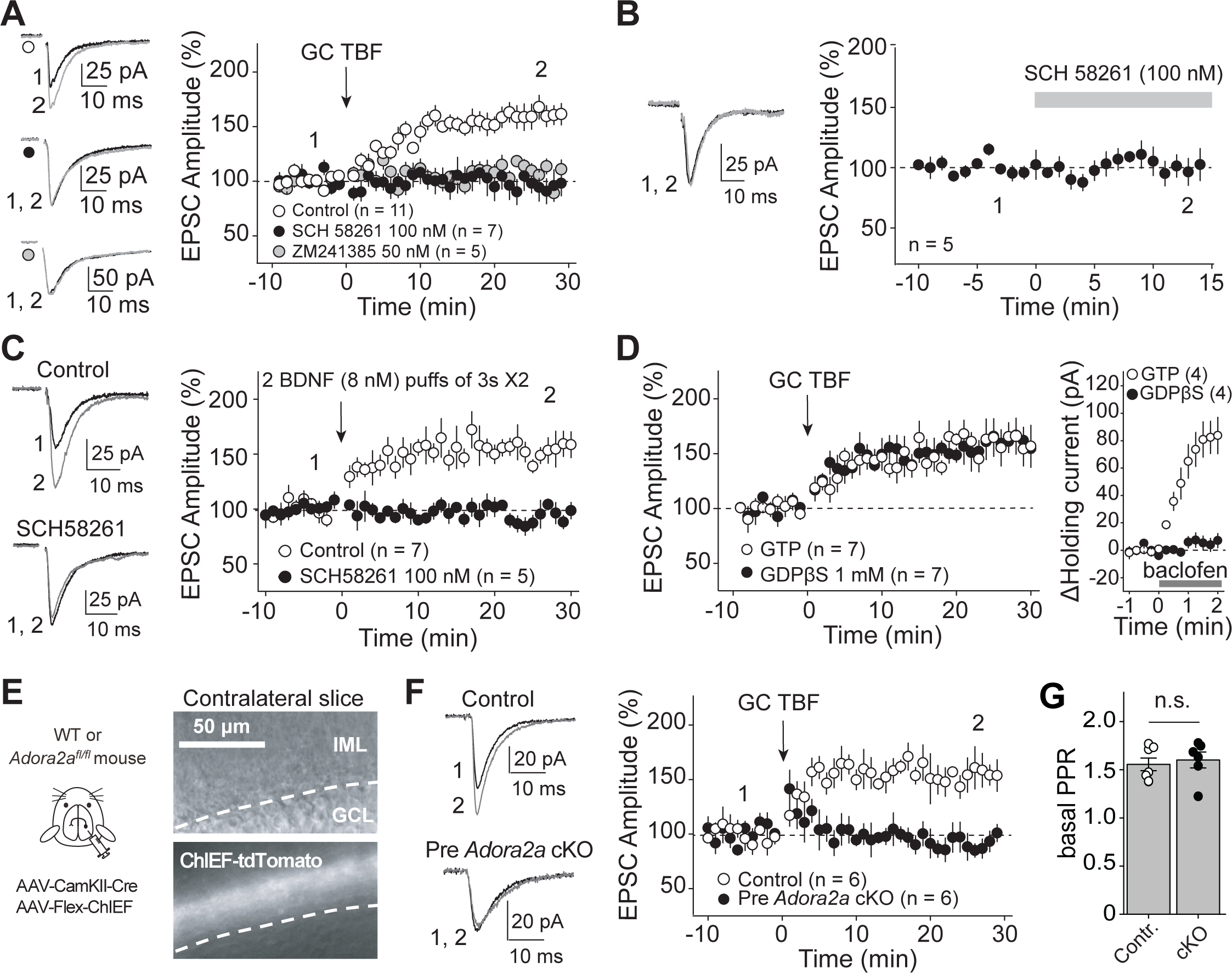
MC-GC LTP requires activation of presynaptic A_2A_Rs. **(A)** Bath application of the A_2A_R selective antagonists SCH 58261 (100 nM) and ZM241385 (50 nM) blocked GC TBF-induced LTP. **(B)** SCH 58261 (100 nM) did not change basal EPSC amplitude. **(C)** SCH 58261 (100 nM) abolished LTP induced by BDNF puffs (8 nM, 2 puffs of 3 s in the IML), as compared with interleaved controls. **(D)** *Left,* loading GDPβS in the recording pipette did not affect TBF-induced LTP. *Right*, interleaved, positive control showing that replacing GTP by 1 mM GDPβS in the internal solution efficiently abolished the GABA_B_ receptor agonist (baclofen, 10 μM)-induced increase in holding current. **(E)** *Left,* A mix of AAV_5_.CamKII.Cre-mCherry and AAV_DJ_.hSyn.Flex.ChIEF.Tdtomato was injected unilaterally into the DG of *Adora2a_fl/fl_* (cKO) or WT (control) mice. *Right,* Infrared/differential interference contrast (IR/DIC, top) and fluorescence (bottom) images show that ChIEF-TdTomato was selectively expressed in putative MC axons of contralateral IML. **(F)** Light-evoked MC EPSCs were recorded in contralateral DG. MC-GC LTP was abolished in presynaptic *Adora2a* conditional knockout mice as compared with controls. **(G)** Basal PPR was similar in control and presynaptic *Adora2a* cKO animals. n.s. p > 0.05., Numbers in parentheses represent number of cells. Data are presented as mean ± SEM.

### Activation of presynaptic A_2A_Rs is sufficient to induce MC-GC LTP

We next tested whether A_2A_R activation was sufficient to induce LTP at MC-GC synapses. Bath application of the A_2A_R agonist CGS21680 (50 nM, 15 min) selectively enhanced MC but not MPP EPSC amplitude (**Figure 3A**). This synapse-specific potentiation was associated with a significant decrease in both PPR and CV (**Figure 3B**), suggesting a presynaptic mechanism of expression, and supporting a presynaptic location of A_2A_Rs. In addition, we tested whether supramammillary (SuM) inputs express functional A_2A_Rs using optogenetic activation of SuM inputs (Tabuchi et al., 2022) (see Methods). Unlike MC EPSCs, SuM EPSCs were not enhanced by CGS21680 bath application (**Figure 3C**). We then directly assessed the contribution of presynaptic A_2A_Rs at MC-GC synapses by testing CGS21680-induced potentiation in presynaptic A_2A_Rs cKO mice. CGS21680 failed to increase MC-GC synaptic transmission in presynaptic *Adora2a* cKO mice (**Figure 3D).** Adding the selective A_2A_R antagonist SCH 58261 (100 nM) during the washout of CGS21680 (50 nM) did not impair the long-lasting potentiation, whereas continuous bath application of SCH 58261 (100 nM) abolished the CGS21680-induced potentiation (**Figure 3E**). These results indicate that activation of presynaptic A_2A_Rs was sufficient to induce LTP at MC-GC synapses.

**Figure 3:**
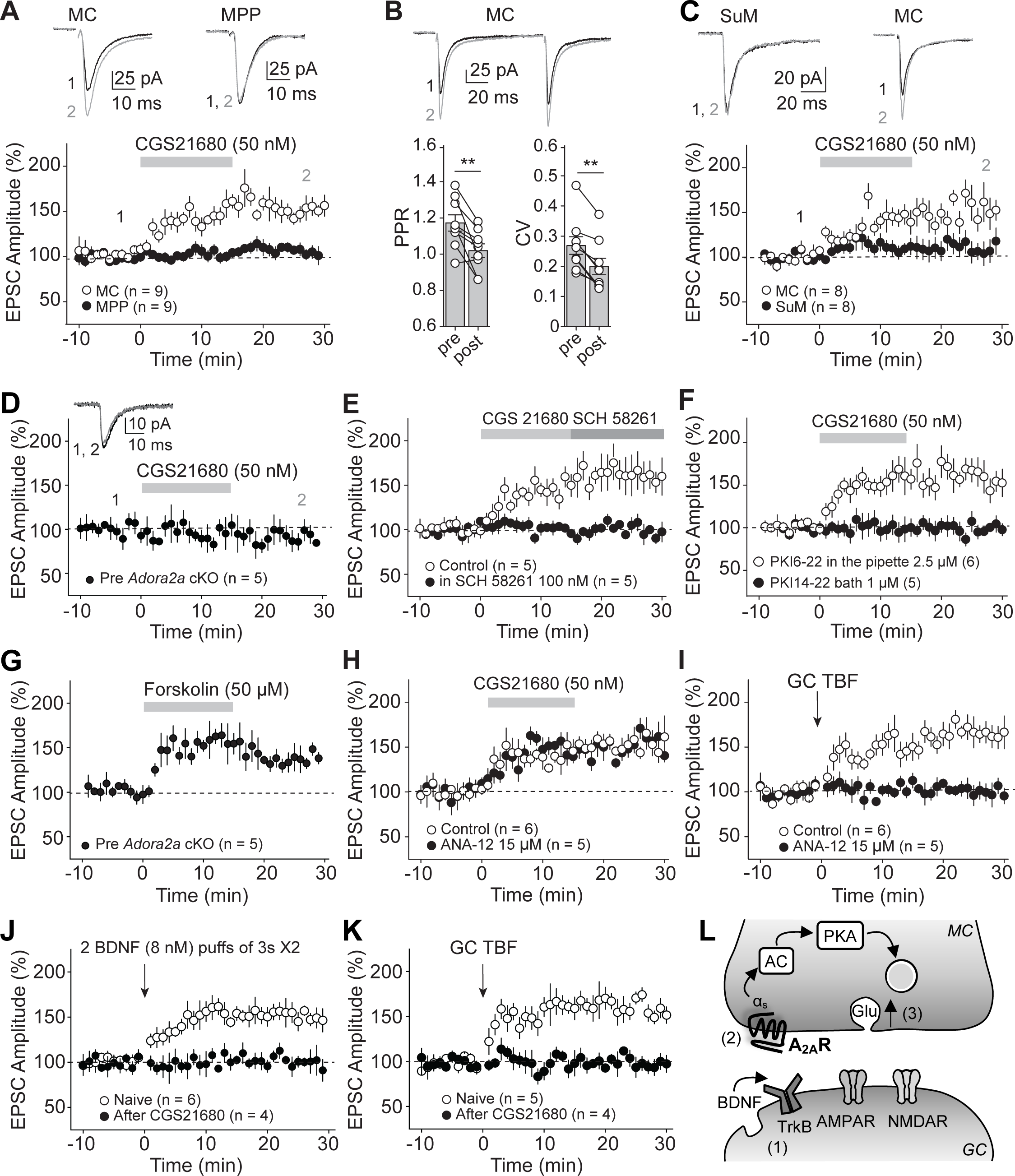
A_2A_R activation is sufficient to trigger PKA-dependent LTP at MC-GC but not at MPP-GC synapses. **(A)** Representative traces *(top)* and time-course summary plot *(bottom)* showing that bath application of the A_2A_R selective agonist CGS21680 (50 nM) potentiated MC-GC but not MPP-GC synaptic transmission. **(B)** CGS21680-induced potentiation at MC-GC synapse was associated with a significant reduction of both PPR and CV. ** p < 0.01; n = 9 cells. **(C)** AAV1-EF1a-DIO-hChR2(H134R)-eYFP was injected into the SuM of VGluT2-Cre mice. Light-evoked SuM EPSCs and electrically-triggered MC EPSCs were monitored in GCs. CGS21680 (50 nM) potentiated MC-GC but not SuM-GC synaptic transmission. **(D)** CGS21680-induced potentiation was abolished in presynaptic *Adora2a* cKO mice. A mix of AAV_5_.CamKII.Cre-mCherry and AAV_DJ_.hSyn.Flex.ChIEF.TdTomato was injected unilaterally into the DG of *Adora2a_fl/fl_*. Light-evoked MC EPSCs were recorded in contralateral DG. **(E)** CGS21680 induced long-lasting potentiation even when the A_2A_R selective antagonist SCH 58261 (100 nM) was included during CGS21680 washout. CGS21680-induced LTP was completely abolished in continuous presence of SCH 58261. **(F)** Loading the selective PKA blocker PKI_6-22_ (2.5 μM) in GCs via the recording pipette did not impair CGS21680-induced LTP while bath application of the cell-permeable PKA inhibitor PKI_14-22_ myristoylated (1 μM) completely blocked LTP. **(G)** Light-evoked MC EPSCs showing that bath application of the adenylyl cyclase activator forskolin (50 μM) induced LTP in presynaptic *Adora2a* cKO mice. **(H)** Bath application of the TrkB antagonist ANA-12 (15 μM) did not impair CGS21680-induced LTP. **(I)** Positive control in interleaved experiments showing that ANA-12 (15 μM) efficiently blocked GC TBF-induced LTP. **(J, K)** Bath application of CGS21680 (50 nM, 15 min) occluded LTP induced with both BDNF (8 nM, 2 puffs of 3 s in the IML, J) and GC TBF (K). **(L)** Cartoon illustrating how activation of presynaptic A_2A_Rs induces PKA-dependent long-lasting increase in glutamate (Glu) release. The presynaptic A_2A_R/PKA pathway is engaged downstream of postsynaptic BDNF/TrkB signaling during LTP induction. Numbers in parentheses represent number of cells. Data are presented as mean ± SEM.

Because the A_2A_R is a G_s_-coupled receptor, we next tested whether CGS21680-induced LTP was PKA-dependent. Bath application of the selective, cell-permeable PKA inhibitor myristoylated PKI_14-22_ (1 μM) (Harris et al., 1997) abolished CGS21680-induced LTP, whereas loading the membrane-impermeable PKA inhibitor PKI_6-22_ (2.5 μM) in GCs via the recording pipette (Hashimotodani *et al*., 2017; Skeberdis et al., 2006) had no effect (**Figure 3F**), suggesting that presynaptic but not postsynaptic PKA activity is required for the A_2A_R agonist-mediated strengthening of MC-GC synaptic transmission. While deleting A_2A_Rs from MCs blocked both GC TBF-LTP (**Figure 2E and 2F**) and CGS21680-LTP (**Figure 3D**), we found that forskolin induced normal LTP in presynaptic *Adora2a* cKO mice (**Figure 3G**). Altogether, these findings indicate that presynaptic A_2A_R activation enhances glutamate release via presynaptic PKA activation, consistent with previous results showing that PKA activity is necessary and sufficient for MC-GC LTP (**Figure 1E-1G, Figure S1J**) (Hashimotodani *et al*., 2017).

Our results thus far indicate that both BDNF/TrkB and adenosine/A_2A_R signaling are involved in MC-GC LTP, and previous work demonstrated that A_2A_Rs facilitate BDNF signaling at some synapses (Sebastiao and Ribeiro, 2009b). We therefore sought to determine potential interactions between these signaling cascades at the MC-GC synapse. The TrkB antagonist ANA-12 did not affect CGS21680-induced LTP (**Figure 3H**), whereas it blocked TBF-LTP in interleaved experiments (**Figure 3I**; see also **Figure S1B**). Lastly, we found that pre-application of CGS21680 (50 nM, 15 min) occluded both BDNF puff-induced and GC TBF-induced LTP (**Figure 3J and 3K**), strongly suggesting that A_2A_R agonism, BDNF, and GC TBF strengthen MC-GC synapses via a common mechanism. These findings, together with our previous results (**Figure 2**), indicate that presynaptic adenosine/A_2A_R signaling mediates LTP downstream of BDNF/TrkB signaling (**Figure 3L**).

### Activation of presynaptic A_1_Rs dampens MC-GC LTP

So far, our data indicate that the release of endogenous adenosine induces MC-GC LTP by activating presynaptic A_2A_Rs (**Figures 2 and 3**). In addition to activating presynaptic A_2A_Rs, adenosine could also activate presynaptic type 1 adenosine receptors (A_1_Rs), G_i/o_-coupled receptors known to suppress neurotransmitter release. To test the role of A_1_Rs in MC-GC plasticity, we first used the selective A_1_R antagonist DPCPX (100 nM). DPCPX increased MC-GC LTP magnitude (**Figure S6A**) but did not affect basal transmission (**Figure S6B**), indicating that adenosine acts on A_1_Rs to dampen MC-GC LTP induction, but A_1_Rs are not tonically activated by adenosine. In addition, we considered that A_1_Rs could mediate presynaptic LTD (Atwood et al., 2014). However, bath application of the selective A_1_R agonist CCPA (50 nM, 15 min) reduced both MC and MPP-mediated transmission reversibly as the reduction was washed out with the A_1_R antagonist DPCPX (100 nM) (**Figure S6C**). This transient A_1_R-induced depression was associated with a reversible increase in both PPR and CV (**Figures S6D and S6E**), suggesting a presynaptic mechanism. Of note, A_2A_R antagonism blocked TBF-LTP induction, but MC-GC transmission was not reduced in a long-term manner (**Figure 2A**), indicating that the net effect of adenosine released during GC TBF is to potentiate MC-GC transmission through A_2A_R activation. Individual MC boutons could contain both A_1_ and A_2A_ receptors. To test this possibility, we examined whether responses elicited by minimal stimulation, a manipulation known to activate one to three MC-GC synapses impinging on GCs (Hashimotodani *et al*., 2017) were sensitive to both A_2A_R and A_1_R agonists. We found that the A_2A_R agonist CGS21680 (50 nM) potentiated the synaptic responses evoked by minimal stimulation of MC axons, whereas subsequent application of the A_1_R agonist CCPA (50 nM) significantly reduced these potentiated responses recorded from the same GC (**Figure S6F and S6G**). These results strongly suggest that functional A_1_ and A_2A_ receptors colocalize in the same MC axon boutons.

#### Anatomical evidence for A_2A_ and A_1_ receptors in MC axon terminals

To test whether MC terminals express A_1_ and A_2A_ receptors, we performed immunoelectron microscopy. Immunoparticles for A_1_Rs were mainly found presynaptically in the molecular layer of the dentate gyrus at MC-GC and perforant path (PP)-GC synapses (**Figures 4A, 4B 4C and 4D**). Remarkably, using an anti-A_2A_R antibody previously validated in A_2A_R-deficient mice (Quiroz et al., 2009), we found strong A_2A_R expression in the presynaptic membrane of asymmetric (presumably glutamatergic) synapses and very few postsynaptic particles in the IML (**Figures 4E, 4F and 4H**). In contrast, a quasi-exclusive postsynaptic expression was detected at asymmetric PP-GC synapses (**Figures 4G, 4I and 4H**). Given the strong projection of MC axons in the IML (Buckmaster et al., 1996), the presynaptic A_2A_R labeling most likely arises from MC axon boutons. While the antibodies were not validated in the DG using immunoelectron microscopy, these anatomical results are entirely consistent with our functional findings. Although both A_1_ and A_2A_ receptors are expressed at MC axon terminals, only the latter engages long-lasting synaptic plasticity.

**Figure 4:**
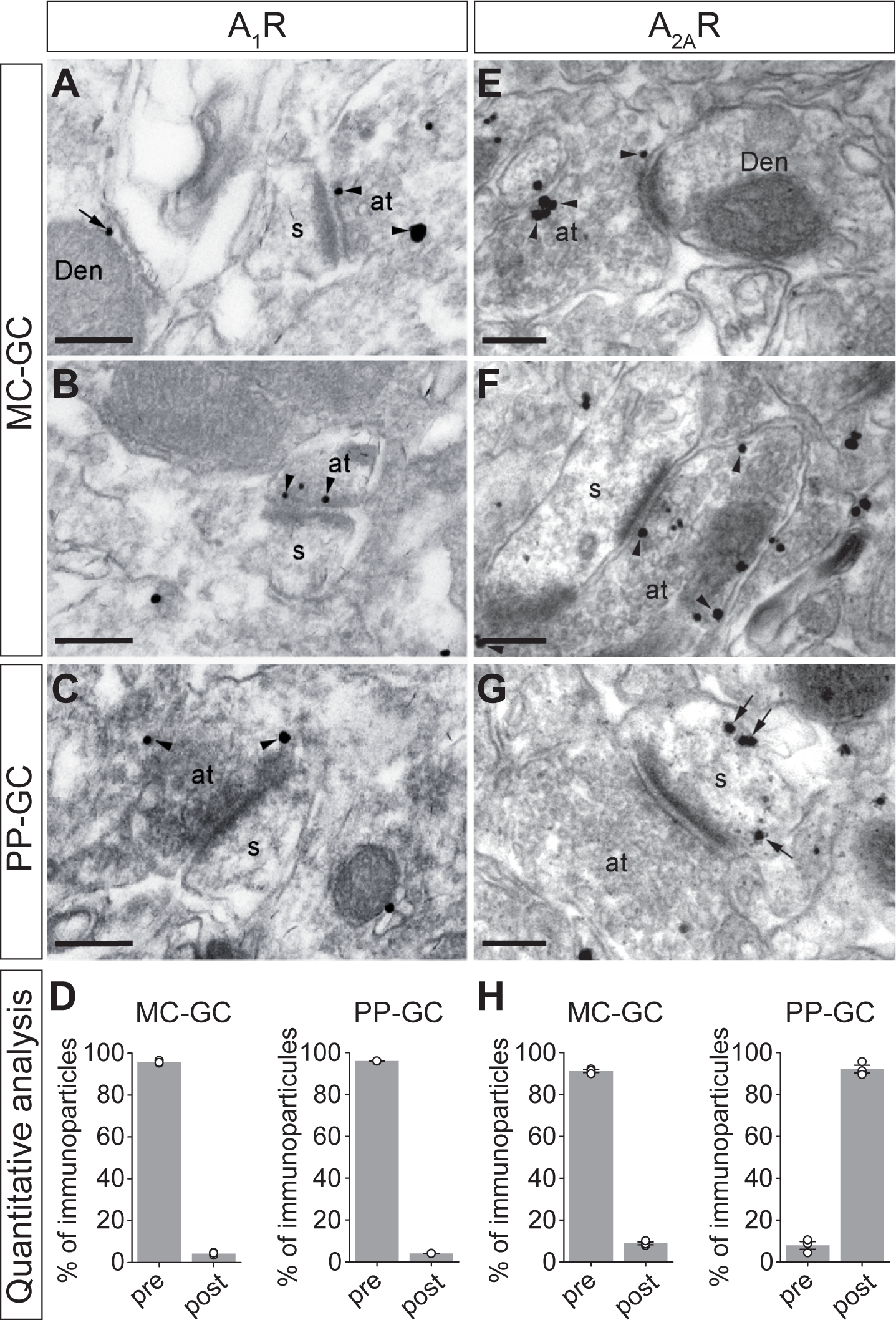
Subcellular localization of A_1_ and A_2A_ receptors in the molecular layer of the dentate gyrus. **(A-C, E-G)** Electron micrographs of the molecular layer of the dentate gyrus showing immunoreactivity for A_1_Rs and A_2A_Rs revealed by pre-embedding immunogold methods. **(A-C)** Both at MC-GC **(A, B)** and PP-GC **(C)** putative synapses, immunoparticles for A_1_R were mainly observed on the presynaptic plasma membrane (arrowheads) of axon terminals (at), with very low frequency in postsynaptic sites (arrows) of spines (s) or dendritic shafts (Den). **(D)** Quantitative analysis of the relative number of immunoparticles found in the presynaptic membrane for A_1_R at MC-GC and MPP-GC putative synapses. From the total number of immunoparticles detected (n = 490 for MC-GC; n = 419 for MPP-GC synapses, N = 3 mice), 468 (95.5%) and 402 (95.9%) were present in presynaptic sites of MC-GC and MPP-GC putative synapses, respectively. **(E-H)** Electron micrographs **(E-G)** and quantitative analysis **(H)** showing that, at putative MC-GC synapses **(E, F** and **H)**, immunoparticles for A_2A_R were mainly detected presynaptically (91.0%) whereas they were mainly found on the postsynaptic plasma membrane (92.0%) of putative PP-GC synapses **(G, H)**. Total number of immunoparticles: n = 668 for MC-GC; n = 690 for MPP-GC synapses, N = 3 mice. Scale bars: 200 nm. Data are presented as mean ± SEM.

### Passive adenosine release from GC is required for presynaptic MC-GC LTP

We next sought to determine the source of adenosine that triggers A_2A_R-mediated LTP. Adenosine can be released from neurons upon activity (Latini and Pedata, 2001; Lovatt *et al*., 2012; Pons-Bennaceur *et al*., 2019; Wall and Dale, 2013) via equilibrative nucleoside transporters (ENTs) (King *et al*., 2006). If adenosine is the retrograde signal mediating MC-GC LTP, interfering with adenosine release from GCs, by blocking ENTs, should impair LTP. As a first approach, we tested whether blocking ENTs can interfere with LTP induction. Bath application of the ENT inhibitors dipyridamole (20 μM) and NBMPR (10 μM) abolished LTP (**Figure 5A**) but did not affect basal MC-GC synaptic transmission (**Figure 5B**). Because ENTs are also implicated in adenosine reuptake, bath application of the ENT blockers may significantly increase extracellular adenosine levels, which, by activating A_2A_Rs may potentiate MC-GC transmission and occlude LTP. Further, the fact that ENT blockers did not affect basal MC-GC synaptic transmission (**Figure 5B**) could be due to the simultaneous activation of A_1_R, which might mask any potential A_2A_R-mediated potentiation. To test this possibility, we bath applied both the ENT blockers and DPCPX (100 nM) to prevent a potential A_1_R-mediated depression of MC-GC synaptic transmission. We found that co-application of the ENT blockers and DPCPX (100 nM) increased MC-GC transmission, and this effect was abolished in the presence of the A_2A_R antagonist SCH 58261 (100 nM) (**Figure 5C**). These results indicate that endogenous adenosine can activate presynaptic A_2A_Rs to potentiate MC-GC synaptic transmission.

**Figure 5:**
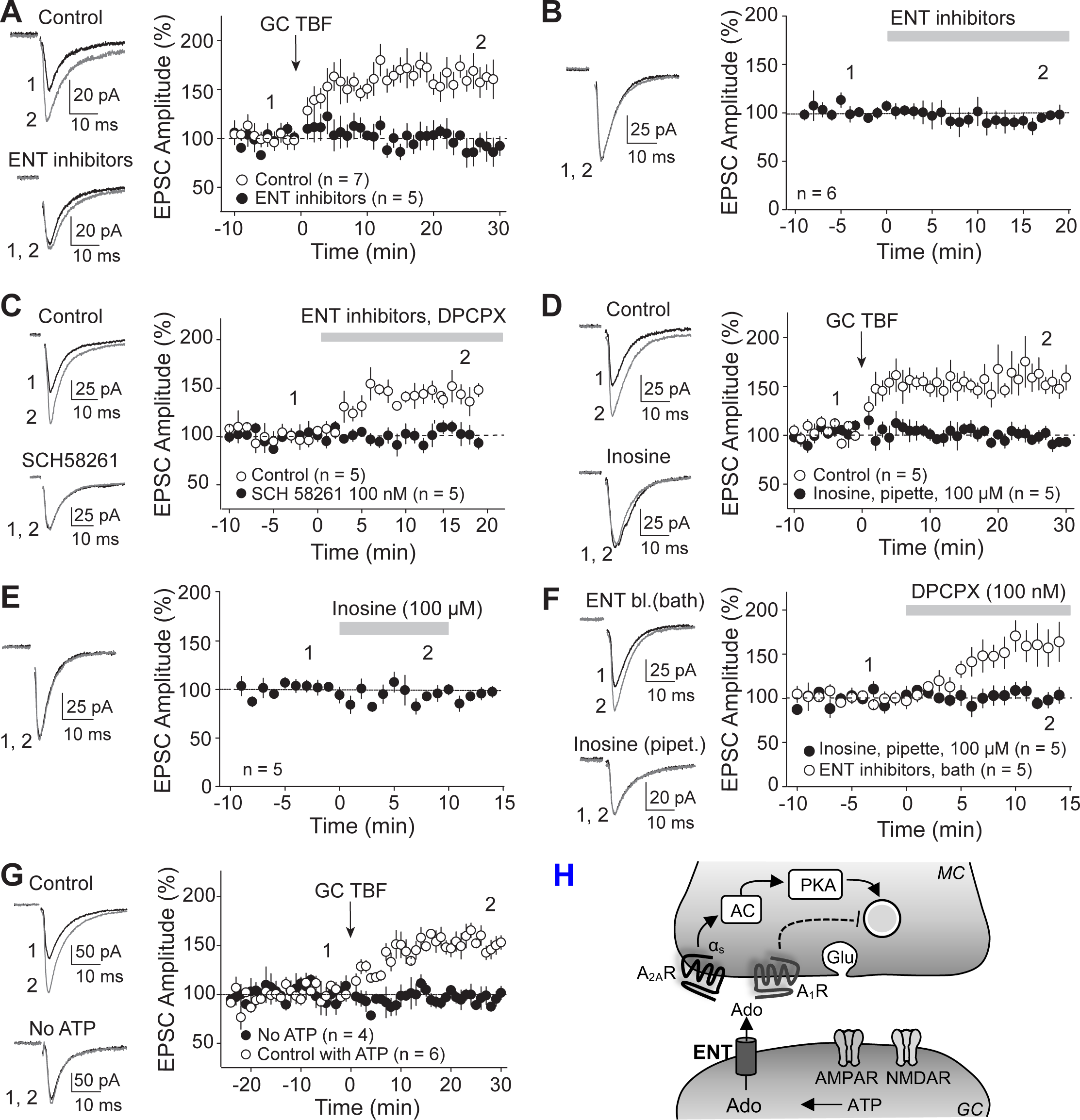
MC-GC LTP requires passive release of adenosine from GCs, via ENTs. **(A)** Representative traces *(left)* and time-course summary plot *(right)* showing that GC TBF failed to induce LTP in presence of the ENT blockers (20 μM of dipyridamole and 10 μM of NBMPR) as compared with interleaved controls. **(B)** Bath application of the ENT blockers (20 μM of dipyridamole and 10 μM of NBMPR) did not change basal MC-GC EPSC amplitude. **(C)** Co-application of the ENT blockers (20 μM of dipyridamole and 10 μM of NBMPR) and the A_1_R antagonist DPCPX (100 nM) increased EPSC amplitude in the control condition but not in presence of the A_2A_R antagonist SCH 58261 (100 nM). **(D)** Intracellular loading of inosine (100 µM) via the patch pipette abolished GC TBF-induced LTP. **(E)** Time course summary plot (right) and representative traces (left) showing that bath application of inosine (100 µM) did not affect basal EPSC amplitude. **(F)** Bath application of DPCPX (100 nM) increased EPSC amplitude when the ENT blockers (20 μM of dipyridamole and 10 μM of NBMPR) were included in he bath but not when inosine (100 µM) was loaded in the postsynaptic neuron via the patch pipette. **(G)** GC TBF failed to induce LTP when ATP was removed from the recording solution as compared to controls. To allow for complete intracellular dialysis, LTP induction protocol was applied 25-35 min after break-in. **(H)** Scheme summarizing the findings. Adenosine is passively released from GCs via ENTs. Adenosine then activates presynaptic A_1_Rs and A_2A_Rs. A_2A_R activation induces a long-lasting PKA-dependent increase in glutamate (Glu) release, whereas A_1_R activation dampens synaptic transmission and LTP. Numbers in parentheses represent number of cells. Data are represented as mean ± SEM.

To directly test whether postsynaptic adenosine release is required for MC-GC LTP, we employed a single-cell approach and selectively blocked ENTs in a single GC. Like previous studies (Lovatt *et al*., 2012; Pons-Bennaceur *et al*., 2019), we loaded inosine (100 µM), a competitive blocker of adenosine efflux through ENTs (King *et al*., 2006), intracellularly *via* the recording pipette. We found that intracellular inosine abolished TBF-LTP (**Figure 5D**), whereas bath application of 100 µM inosine had no effect on MC-GC synaptic transmission (**Figure 5E**). Bath application of DPCPX increased MC EPSC amplitude when ENT inhibitors were continuously bath applied but not when inosine (100 µM) was loaded in a single GC (**Figure 5F**), confirming that intracellular inosine did not increase adenosine tone. Intracellular accumulation of adenosine can result from sequential dephosphorylation of ATP (Arch and Newsholme, 1978). Consistent with this possibility, removing ATP from the internal recording solution abolished LTP (**Figure 5G**). Altogether these findings indicate that adenosine, passively released from GC through ENTs, acts as a retrograde messenger at MC-GC synapses (**Figure 5H**).

### Repetitive neuronal activity induces adenosine release via a BDNF/TrkB-dependent mechanism

To visualize adenosine release during MC-GC LTP induction, we utilized the genetically encoded sensor for adenosine GRAB_Ado1.0m_ (Peng et al., 2020; Wu et al., 2023), which was selectively expressed in commissural MC axons (see Methods) and generated a clear fluorescence signal in the contralateral IML (**Figures 6A, 6B and 6D**). The MC burst stimulation protocol that triggers synaptically-induced MC-GC LTP in acute hippocampal slices also triggered a robust, albeit transient increase in the GRAB_Ado1.0m_ signal. As expected (Peng *et al*., 2020), this enhancement was abolished when the MC BS protocol was delivered in the presence of the A_2A_R antagonist SCH58261 (**Figures 6B, 6C and 6F**). Consistent with A_2A_R signaling being engaged downstream of TrkB activation, the burst-induced increase of the GRAB_Ado1.0m_-mediated fluorescence was abolished in the presence of the TrkB antagonist ANA-12 (15 µM), and in postsynaptic *TrkB* cKO and *Bdnf* cKO mice (**Figures 6D-6F**). Altogether these results demonstrate that MC repetitive activity releases adenosine in a postsynaptic BDNF/TrkB-dependent manner.

**Figure 6:**
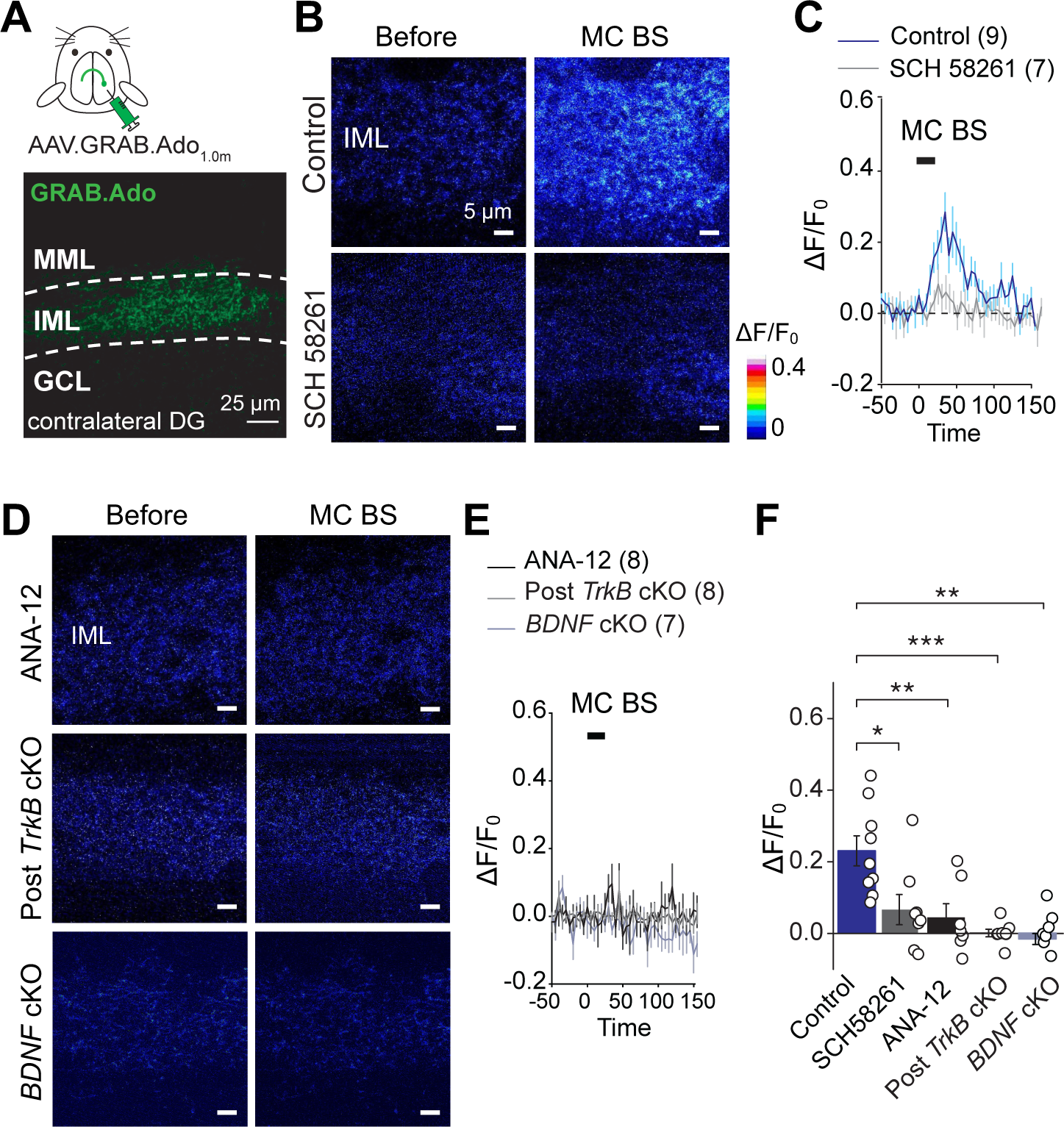
Induction of MC-GC LTP triggers a transient TrkB-dependent increase in extracellular adenosine level. **(A)** AAV_9_.hSyn.GRAB.Ado1.0m (GRAB_Ado_) was injected unilaterally in the DG of WT mice (*top*). Two-photon image (*bottom*) showing GRAB_Ado_ was selectively expressed in putative MC axons in contralateral IML. **(B, C)** Two-photon images of the IML **(B)** showing GRAB_Ado_ fluorescence intensity increased during burst electrical stimulation of MC axon terminals (MC BS) in normal ACSF (control) but not in continuous presence of the A_2A_ receptor antagonist SCH 58261 (100 nM). Time-course summary plot of the average fractional fluorescence changes (ΔF/F_0_) with time are shown in **C** **(D, E)** Two-photon images of the IML **(D)** and time-course summary plot **(E)** showing how MC BS failed to increase GRAB_Ado_ fluorescence intensity in the continuous presence of the TrkB antagonist ANA-12 (15 µM) and when TrkB (Post *TrkB* cKO) or BDNF (Post *Bdnf* cKO) was conditionally knocked out from GCs. **(F)** Quantification of the averaged responses during burst stimulation of MCs (15-25 s) * p < 0.05, ** p < 0.01, *** p < 0.001; one-way ANOVA. Number of slices are shown between parentheses. Data are represented as mean ± SEM.

### *In vivo* activation of A_2A_Rs facilitates seizure occurrence

We recently found that MC-GC LTP can be triggered by seizure activity induced with kainic acid (KA) intraperitoneal (i.p.) administration and that this LTP further promotes seizures (Nasrallah *et al*., 2022). Given our new findings that neuronal activity releases adenosine via TrkB to induce A_2A_R-dependent MC-GC LTP, we hypothesized that interfering with adenosine/A_2A_R signaling should prevent seizure-induced LTP and reduce KA-induced seizures. Moreover, seizure activity should release adenosine in a TrkB-dependent manner. As expected for a presynaptic form of plasticity, seizure-induced LTP increases neurotransmitter release as indicated by a decrease in PPR (Nasrallah *et al*., 2022). While *Adora2a* deletion from MC axons did not alter basal PPR (**Figure 2G**), it prevented the KA-induced decrease in PPR (**Figure 7A and 7B**). These results indicate that presynaptic A_2A_Rs mediate seizure-induced LTP at MC-GC synapses. Moreover, *Adora2a* deletion increased latency to KA-induced convulsive seizures (**Figures 7C-7F**), and *TrkB* deletion increased latency to convulsive seizures and reduced sum seizure score (**Figures 7G and 7H**). These results strongly suggest that both A2ARs and TrkB are proconvulsant, consistent with the seizure-induced release of adenosine in a TrkB-dependent manner.

**Figure 7:**
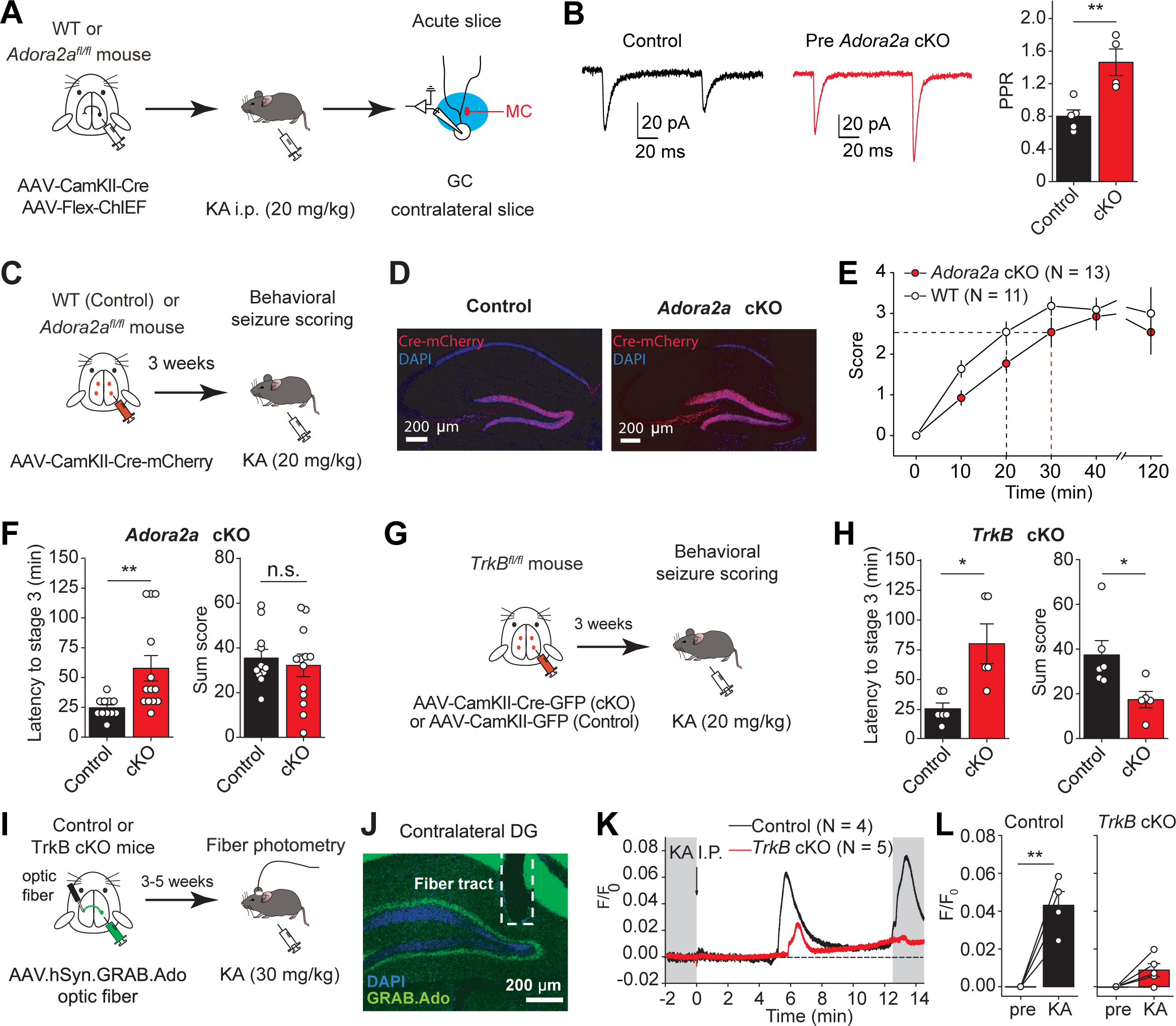
*In vivo* release of adenosine during acute seizures plays a pro-convulsant role by activating A_2A_Rs. **(A)** Experimental timeline. A mix of AAV_5_.CamKII.Cre-mCherry and AAV_DJ_.hSyn.Flex.ChIEF.TdTomato was injected unilaterally into the DG of *Adora2a^fl/fl^* (*Adora2a* cKO) or WT (control) mice. Seizures were induced using a single KA i.p. (20 mg/kg) injection and acute hippocampal slices were prepared 25 min after KA injection. Whole-cell recordings were performed from GC in the contralateral DG and MC-GC light-evoked synaptic responses were monitored. **(B)** KA-induced seizures decreased PPR in control but not in presynaptic *Adora2a* cKO conditions. ** p < 0.01. **(C)** AAV_5_-CaMKII-Cre-mCherry was injected bilaterally into ventral and dorsal DG of WT (control) and *Adora2a_fl/fl_* (cKO) mice. Mouse behavior was assessed for 120 min following KA (20 mg/kg i.p.) administration. **(D)** Confocal images showing the viral expression in the DG of WT (control) and *Adora2a_fl/fl_* (*Adora2a* cKO) mice. **(E, F)** Deletion of *Adora2a* from DG excitatory neurons (*Adora2a_fl/fl_* mice injected with AAV_5_-CaMKII-Cre-mCherry) significantly increased latency to convulsive seizures **(E, F),** but did not affect seizure severity as compared to control animals. Note the shift in seizure score time course toward the right **(E)**. ** p < 0.01, n.s. p > 0.05. **(G, H)** AAV-CaMKII-eGFP (control) or AAV-CaMKII-Cre-GFP (cKO) was injected bilaterally into ventral and dorsal DG of *TrkB_fl/fl_* mice. Behavioral seizures were monitored and scored for 120 min **(G)**. Deletion of *TrkB* from hippocampal excitatory neurons induced significant increase in latency to convulsive seizures **(E)** and a decrease in sum score **(H)** and as compared with controls. * p < 0.05. **(I)** Experimental timeline. AAV_9_.hSyn.GRAB.Ado1.0m (GRAB_Ado_) was injected unilaterally in the DG of *TrkB_fl/fl_* mice. In the contralateral DG, both control (AAV-CaMKII-mCherry) or Cre-expressing AAV (AAV-CaMKII-Cre-mCherry, TrkB cKO) was injected and an optic fiber was implanted above the contralateral IML. Fiber photometry was performed 3-5 weeks later to assess GRAB_Ado_ fluorescence intensity before and after acute seizure induction with kainic acid (KA, 30 mg/kg i.p.), in control and *TrkB* cKO mice. **(J)** Confocal image showing fiber tract and GRAB_Ado_ expression in the contralateral IML. **(K, L)** Time-course of a control and a *TrkB* cKO representative experiments **(K)** and summary histogram **(L)** showing how KA (30 mg/kg i.p.) administration increased the average fractional fluorescence (ΔF/F_0_) of GRAB_Ado_ in control mice, an effect that was significantly reduced in *TrkB* cKO animals. ** p < 0.01, n.s. p > 0.05. Numbers in parentheses represent number of mice. Data are presented as mean ± SEM.

Using *in vivo* two-photon imaging in awake mice, we recently reported that KA-induced seizures trigger a massive increase in both MC and GC activity (Nasrallah *et al*., 2022). To test whether seizure activity triggers a TrkB-dependent release of adenosine *in vivo*, we expressed GRAB_Ado1.0m_ in commissural MC axons and recorded the fluorescence signals using fiber photometry in freely moving mice (**Figures 7I and 7J**). We found a large increase in GRAB_Ado1.0m_ fluorescence following i.p. injection of KA, and this fluorescence signal was significantly reduced in *TrkB* cKO mice (**Figures 7K and 7L**). Of note, both control and *TrkB* cKO mice reached stage 3 convulsive seizures, discarding potential failure of seizure induction. Altogether, these results not only demonstrate the TrkB-mediated *in vivo* release of adenosine following initial seizures, but also strongly suggest that adenosine signaling may promote seizures via A_2A_Rs.

## DISCUSSION

In this study, we discovered a novel retrograde signaling mechanism that mediates presynaptic strengthening within a critical hippocampal circuit (**Figure S7**). Specifically, repetitive firing of a single postsynaptic neuron was sufficient to trigger presynaptic LTP at MC-GC but not neighboring MPP synapses. We demonstrated that functional A_2A_Rs are expressed at MC axon terminals but not at MPP axon terminals and that A_2A_R activation is necessary and sufficient to induce MC-GC LTP. During LTP induction GCs passively release adenosine through ENTs in a postsynaptic TrkB- and activity-dependent manner. Thus, our findings establish adenosine/A_2A_R as a novel retrograde signaling mechanism. Remarkably, our results uncovered that TrkB can signal through adenosine. Moreover, we found that adenosine is released *in vivo* during initial seizures in a TrkB-dependent manner and both *Adora2a* and *Trkb* cKO have anti-convulsant effects.

### GC TBF induces presynaptic LTP at MC-GC synapses

The present study revealed that direct activation of a single GC with a physiologically relevant pattern of activity (Danielson et al., 2017; Diamantaki et al., 2016; GoodSmith et al., 2017; Henze et al., 2002; Pernia-Andrade and Jonas, 2014; Senzai and Buzsaki, 2017) was sufficient to induce presynaptic LTP at MC-GC synapses without changing the strength of MPP neighboring inputs. Although the activity of a single GC could have induced LTP by recruiting the recurrent GC-MC-GC circuit, the following two observations are against this possibility. First, GC TBF induced normal LTP even in the continuous presence of the selective group II mGluR agonist DCG IV, which abolishes GC-MC (Lysetskiy *et al*., 2005) but not MC-GC transmission (Chiu and Castillo, 2008). Second, TBF-LTP was normally induced at optogenetically-activated synaptic inputs arising from contralateral MC axons that are cut off from their cell bodies in hippocampal slices. Importantly, TBF-LTP and synaptically-induced LTP likely share a common mechanism (**Figure S2**). Like synaptically-induced MC-GC LTP, which requires BDNF release from GCs (Berthoux *et al*., 2023), we now report that TBF-LTP requires postsynaptic calcium rise and BDNF release via SNARE-dependent exocytosis (Leschik et al., 2019; Shimojo *et al*., 2015). While synaptically-induced LTP is an input-specific phenomenon (Hashimotodani *et al*., 2017), TBF-induced LTP is not, likely reflecting the dissimilar induction protocols. Presynaptic burst stimulation releases adenosine from the postsynaptic compartment of activated synapses only. In contrast, TBF is expected to release adenosine from multiple synapses due to action potential-driven release of BDNF from GCs and autocrine activation of postsynaptic TrkB (Berthoux *et al*., 2023).

### Adenosine is released from GCs in an activity and TrkB-dependent manner

Using the genetically encoded adenosine sensor GRAB_Ado1.0m_ (Peng *et al*., 2020; Wu *et al*., 2023), we detected a phasic increase in extracellular adenosine following both MC-GC LTP induction *ex vivo* and during seizure activity *in vivo*. Importantly, adenosine release required intact BDNF/TrkB signaling (**Figure 6D-6F**, **Figure 7K and 7L**). Previous studies have shown that adenosine can be released from neurons upon activity via ENTs (King *et al*., 2006; Latini and Pedata, 2001; Lovatt *et al*., 2012; Pons-Bennaceur *et al*., 2019; Wall and Dale, 2013; Wu *et al*., 2023). In support of this mechanism, we found that interfering with ENT-mediated release of adenosine from a single postsynaptic GC abolished MC-GC LTP, or not including ATP in the recording pipette, indicating that the passive release of adenosine from GCs is essential for this form of plasticity. Adenosine arising from ATP extracellular conversion (Ribeiro and Sebastiao, 1987; Zimmermann, 2006) or other cell types such as interneurons and glial cells (Latini and Pedata, 2001; Wall and Dale, 2008) could also strengthen MC-GC synaptic transmission.

Intracellular accumulation of adenosine can be the consequence of sequential dephosphorylation of ATP or the hydrolysis of S-adenosylho-mocysteine (Arch and Newsholme, 1978). During MC-GC LTP induction, formation of intracellular adenosine in GCs likely results from robust postsynaptic TrkB activation and sequential dephosphorylation of ATP through its kinase activity. In support of this scenario are the following observations. First, postsynaptic ATP is required for MC-GC LTP (**Figure 5F**). Second, MC-GC LTP induction releases BDNF that activates postsynaptic TrkB on GCs (Berthoux *et al*., 2023; Hashimotodani *et al*., 2017). Third, activity-induced adenosine release in the IML is postsynaptic BDNF/TrkB-dependent, as it is abolished in the presence of the TrkB selective antagonist ANA-12 and by genetically removing *Bdnf* and *TrkB* from GCs (**Figure 6D-6F and Figure 7I-L**). Fourth, adenosine/A_2A_R signaling is engaged downstream of TrkB activation given that BDNF-induced LTP was abolished by A_2A_R antagonism (**Figure 2C**), whereas TrkB antagonism did not affect A_2A_R agonist-induced LTP (**Figure 3E**). Lastly, while BDNF activates postsynaptic TrkB (Berthoux *et al*., 2023; Hashimotodani *et al*., 2017) and adenosine acts via presynaptic A_2A_Rs to induce MC-GC LTP, we cannot discard physical interaction or transactivation, a process whereby A_2A_R activation can induce TrkB phosphorylation in the absence of neurotrophins (Lee and Chao, 2001).

To the best of our knowledge, we provided the first direct evidence that adenosine release can occur in a TrkB-dependent manner. Such a mechanism could explain, at least in part, BDNF and adenosine signaling interactions reported at other synapses in the brain. Previous studies in the CA1 area have shown that A_2A_Rs can affect synaptic transmission and plasticity by modulating BDNF-TrkB signaling (Diogenes *et al*., 2004; Fontinha *et al*., 2008; Rodrigues et al., 2014). Such modulation has recently been implicated in adult neurogenesis (Ribeiro et al., 2021). While these studies suggest that A_2A_Rs act upstream of BDNF, our findings demonstrate that BDNF/TrkB signaling can also act upstream A_2A_Rs by promoting adenosine release. Consistent with our findings at the hippocampal MC-GC synapse, BDNF-induced facilitation at the neuromuscular junction was abolished in presence of A_2A_R antagonists but also by PKA inhibitors, while A_2A_R agonist-induced increase in neurotransmitter release is TrkB-independent (Correia-de-Sa et al., 1991; Pousinha et al., 2006; Sebastiao and Ribeiro, 2009b). In conclusion, our findings support a model whereby neuronal activity releases adenosine in a TrkB-dependent manner, and therefore, adenosine release emerges as a non-canonical TrkB signaling mechanism. Further work is required to determine the generalizability of this model in other brain areas.

### Retrograde adenosine/A_2A_R signaling mediates presynaptic LTP

Retrograde signaling is a common mechanism in presynaptic forms of long-term plasticity (Castillo, 2012; Monday *et al*., 2018). While several retrograde signals have been identified throughout the brain (Regehr *et al*., 2009), here we uncovered adenosine/A_2A_R as a novel form of retrograde signaling. Single cell manipulations revealed that interfering with adenosine release from a single postsynaptic neuron abolished MC-GC LTP, induced by GC firing alone. Furthermore, by combining pharmacology, presynaptic *Adora2a* deletion using cKO mice, and immunoelectron microscopy, we demonstrated that activation of presynaptic A_2A_Rs was necessary and sufficient for MC-GC LTP. Despite the relatively low expression levels of A_2A_R in the hippocampus (Dixon et al., 1996), previous work has demonstrated that A_2A_R signaling can regulate hippocampal synaptic plasticity (Dias et al., 2013; Duster et al., 2014). To our knowledge, our study not only provides the first evidence of adenosine/A_2A_R as a retrograde signaling system but also the first evidence that this signaling system can mediate presynaptic LTP. As G_s_-coupled receptors, and as supported by our findings, A_2A_Rs on MC axon terminals likely activate the cAMP/PKA cascade, and as for other forms of presynaptic LTP (Castillo, 2012), this activation engages a long-lasting increase of glutamate release whose downstream mechanism is poorly understood. Whether retrograde adenosine/A_2A_R signaling can mediate activity-dependent strengthening at other synapses in the brain, including in the striatum and globus pallidus, where presynaptic A_2A_Rs are highly expressed (Kase, 2001), remains to be investigated.

### Retrograde adenosine/A_1_Rs signaling dampens LTP induction

As previously reported at other synapses (Correia-de-Sa *et al*., 1991; Pousinha et al., 2010), our findings indicate that postsynaptically released adenosine simultaneously activates A_1_Rs and A_2A_Rs localized at MC terminals. A_1_Rs and A_2A_Rs can localize at the same nerve terminal (Correia-de-Sa et al., 1996) and even form heteromers (Sebastiao and Ribeiro, 2009a). While A_1_R agonism transiently inhibited glutamate release, A_2A_R agonism increased glutamate release in a long-lasting manner. Consistent with these pharmacological observations, the net effect mediated by transient release of endogenous adenosine during LTP induction was a long-lasting enhancement of glutamate release. We also found A_1_R-mediated dampening of LTP which has previously been observed at other synapses (Alzheimer et al., 1991; Forghani and Krnjevic, 1995). A possible explanation for this dampening is that presynaptic A_1_Rs, which are G_i/o_-coupled, decrease the level of cAMP, which is required for MC-GC LTP (Hashimotodani *et al*., 2017). Type 1 cannabinoid receptor, another G_i/o_-coupled receptor that is highly expressed at MC axon terminals, also dampens MC-GC LTP (Jensen *et al*., 2021). Thus, the induction of this form of plasticity is tightly controlled by two distinct retrograde signals, adenosine and endocannabinoids, which, by activating presynaptic G_i/o_-coupled receptors, dampen LTP induction. Both retrograde signals may prevent runaway activity of the MC-GC-MC recurrent excitatory circuit.

### Adenosine/A_2A_R retrograde signaling in the dentate gyrus has physiological and pathological relevance

Growing evidence indicates that MCs are critically involved in hippocampal-dependent learning (Scharfman, 2016). For example, silencing MCs selectively impairs the retrieval of spatial memory (Bui *et al*., 2018) and novelty-induced contextual memory acquisition (Fredes et al., 2021). MC-GC LTP is a robust form of presynaptic plasticity that can be elicited *in vivo* upon experience (Berthoux *et al*., 2023) and may contribute significantly to learning and memory by changing information flow in the DG (Hashimotodani *et al*., 2017). While GCs exhibit extremely sparse and selective firing in a single place field (Diamantaki *et al*., 2016), we found that MC-GC LTP can be triggered by GC firing alone, suggesting that this plasticity could be implicated in experience-driven refinement of DG circuitry and memory. Previous work indicates that adenosine/A_2A_R signaling may contribute to hippocampal-dependent memory (Duster *et al*., 2014; Sebastiao and Ribeiro, 2009a). Our findings showing a key role for adenosine/A_2A_R signaling in MC-GC LTP could explain, at least in part, memory impairments found in both A_2A_R antagonist-injected animals (Fontinha et al., 2009) and hippocampal *Adora2a* cKO animals (Wei et al., 2014). Demonstrating the precise role of adenosine/A_2A_R retrograde signaling at MC-GC synapses in memory requires selective manipulation of A_2A_Rs at MC axon terminals using tools that are not currently available.

Adenosinergic signaling is critically involved in epilepsy (Beamer et al., 2021). Global A_2A_R genetic deletion and selective A_2A_R antagonism attenuate both the seizure progression (El Yacoubi et al., 2008; 2009) and seizure-induced neuronal damage (Jones et al., 1998; Lee et al., 2004; Rosim et al., 2011). In contrast, A_2A_R activation lowers the seizure threshold (Fukuda et al., 2011) and increased A_2A_R function promotes epilepsy (Sandau et al., 2016). Although the precise mechanism by which A_2A_Rs participate in temporal lobe epilepsy remains unclear, our current findings provide a potential explanation for the pro-epileptic role of A_2A_Rs, consistent with our previous findings that initial seizures trigger MC-GC LTP and that uncontrolled LTP is sufficient to promote subsequent seizures (Nasrallah *et al*., 2022). The BDNF/TrkB-mediated release of adenosine from GCs and the consequent strengthening of MC-GC synaptic transmission may also explain, at least in part, the pro-epileptic role of BDNF/TrkB signaling (McNamara and Scharfman, 2012). Our current study demonstrating adenosine/A_2A_R-dependent strengthening of MC-GC synapses reveals a novel mechanism whereby MC activity may contribute to early stages of temporal lobe epilepsy.

## Acknowledgments

We thank all Castillo lab members for their invaluable feedback. We thank Dr. Anita Autry (Einstein) for sharing her photometry system and for the technical support provided by her lab members, especially Ilaria Carta and Dr. Giovanni Podda., We thank Subrina Persaud for assisting with confocal image acquisition. We thank Dr. Yulong Li (Peking University) for helpful discussions about the adenosine sensor, Dr. Pascal Kaeser (Harvard University) for sharing an AAV-hSyn-Flex-ChIEF-TdTomato plasmid, and Dr. Lisa Monteggia (Vanderbilt University) for sharing *TrkB^fl/fl^* and *Bdnf^fl/fl^* mice, and Drs. Gaël Barthet and Christophe Mulle, Université de Bordeaux, for sharing the C1ql2 constructs. This work was supported by NIH grants R01 MH116673, R01MH125772, and R01 NS 113600 to P.E.C.. K.N. was partially supported by a Postdoctoral Research Fellowship (American Epilepsy Society), the Fondation pour la Recherche Médicale (postdoctoral fellowship for research abroad), and the Fondation Bettencourt Schueller (Prix pour les Jeunes Chercheurs 2016). C.B. was partially supported by a Postdoctoral Research Fellowship (American Epilepsy Society). Y.H. was supported by grants from JSPS KAKENHI (20H03358, 23H04240, and 23K18167). A.E.C. was partially supported by a Ruth L. Kirschstein Award (F32 NS071821), a NARSAD Young Investigator Grant from the Brain & Behavior Research Foundation, and by Chilean Fondecyt regular (#1201848). M.G. was supported by a Ruth L. Kirschstein Award (F31MH122134). R.L. was funded by the Spanish MCIN/AEI/ 10.13039/501100011033 and by “ERDF A way of making Europe” (PID2021-125875OB-I00) and Junta de Comunidades de Castilla-La Mancha (SBPLY/21/180501/000064). Confocal images were obtained at the Einstein Imaging Core (supported by the Rose F. Kennedy Intellectual Disabilities Research Center - shared instrument grant 1S10OD25295 to Konstantin Dobrenis).

## Author contributions

K.N. and P.E.C. designed studies and wrote the manuscript. K.N. performed experiments and analyzed all data except for the *in vitro* adenosine measurements that were designed, performed, and analyzed by C.B.. R.L. performed and analyzed immunoelectron microscopy experiments. M.G. tested the role of A1 receptors. Y.H and A.C designed, performed, and analyzed early experiments assessing conventional retrograde signals. All authors read and edited the manuscript.

## STAR METHODS

### Experimental Model and Subject Details

P19-P30 Sprague-Dawley rats and P50-P70 C57BL/6, floxed TrkB (*TrkB_fl/fl_*), floxed BDNF (*Bdnf_fl/fl_*), floxed *Adora2a* (*Adora2a_fl/fl_*), and VGluT2-Cre mice (Jackson labs, Slc17a6_tm2(cre)Lowl_/J, stock #016963) mice were used. Males and females were equally used. All animals were group housed in a standard 12 hr light/12 hr dark cycle and had free access to food and water. Animal handling and use followed a protocol approved by the Institutional Animal Care and Use Committee of Albert Einstein College of Medicine, in accordance with the National Institutes of Health guidelines. Animal care and handling for immunoelectron microscopy studies followed a protocol approved by the Institutional Animal Care and Use Committee of Universidad Castilla-La Mancha in accordance with Spanish (RD 1201/2005) and European Union (86/609/EC) regulations. The supramammillary experiments were approved by the Animal Care and Use Committee of Doshisha University and were performed following the committee guidelines. *TrkB_fl/fl_* and *Bdnf_fl/fl_* mice generated by Dr. Luis Parada were kindly donated by Dr. Lisa Monteggia (University of Texas, Southwestern Medical Center). *Adora2_fl/fl_* were obtained from Jackson Laboratory (B6;129-Adora2atm1Dyj/J, Jax 010687).

### Hippocampal slice preparation

Acute transverse hippocampal slices were prepared from Sprague-Dawley rats (400 μm thick) and mice (300-μm thick). Animals were anesthetized with isofluorane and euthanized following institutional regulations. The hippocampi were then removed and cut using a VT1200s microslicer (Leica Microsystems Co.) in a cutting solution. Hippocampal slices from rats were prepared using a cutting solution containing (in mM): 215 sucrose, 2.5 KCl, 26 NaHCO_3_, 1.6 NaH_2_PO_4_, 1 CaCl_2_, 4 MgCl_2_, 4 MgSO_4_ and 20 D-glucose. 30 min post-sectioning, the cutting medium was gradually switched to an extracellular artificial cerebrospinal (ACSF) recording solution containing (in mM): 124 NaCl, 2.5 KCl, 26 NaHCO_3_, 1 NaH_2_PO_4_, 2.5 CaCl_2_, 1.3 MgSO_4_ and 10 D-glucose. Slices were incubated for at least 40 min in the ACSF solution before recording. Mouse hippocampal slices were prepared using an NMDG-based cutting solution containing (in mM): 93 N-Methyl-d-glucamin, 2.5 KCl, 1.25 NaH_2_PO_4_, 30 NaHCO_3_, 20 HEPES, 25 D-glucose, 2 Thiourea, 5 Na-Ascorbate, 3 Na-Pyruvate, 0.5 CaCl_2_, 10 MgCl_2_. These slices were then transferred to 32°C ACSF for 30 min and then kept at room temperature for at least 1h before recording. All solutions were equilibrated with 95% O_2_ and 5% CO_2_ (pH 7.4).

### Electrophysiology

Recordings were performed at 28 ± 1°C, otherwise stated, in a submersion-type recording chamber perfused at 2 mL/min with ACSF supplemented with the GABA_A_ and the GABA_B_ receptor antagonists, picrotoxin (100 μM) and CGP55845 hydrochloride (3 μM), respectively. Whole-cell patch-clamp recordings using a Multiclamp 700A amplifier (Molecular Devices) were obtained from GCs voltage clamped at −60 mV using patch-type pipette electrodes (∼3-4 MΩ) containing a potassium-based internal solution (in mM): 135 KMeSO_4_, 5 KCl, 1 CaCl_2_, 5 NaOH, 10 HEPES, 5 MgATP, 0.4 Na_3_GTP, 5 EGTA and 10 D-glucose, pH 7.2 (288-291 mOsm). For IPSC recordings in CA1, whole-cell voltage clamp recordings were obtained from CA1 pyramidal neurons voltage clamped at V_h_= 10 mV using a cesium-based internal solution containing (in mM): 131 cesium gluconate, 8 NaCl, 1 CaCl_2_, 10 EGTA, 10 D-glucose and 10 HEPES, pH 7.2 (285-290 mOsm). IPSCs were evoked using an electrical stimulating pipette placed in CA1 *stratum radiatum*, and recordings were performed in the continuous presence of ionotropic glutamate receptor antagonists NBQX (10 μM) and D-APV (50 μM). LTD of inhibitory inputs onto CA1 pyramidal neurons (iLTD) was induced using theta burst-stimulation protocol (TBS, 10 bursts at 5 Hz of 5 pulses at 100 Hz, repeated every 5 s, 4 times). Series resistance (∼6-25 MΩ) was monitored throughout all experiments with a −5 mV, 80 ms voltage step, and cells that exhibited a significant change in series resistance (> 15%) were excluded from the analysis. Botox experiments were performed at 32°C. Botox was supplemented with 5 mM dithiothreitol (DTT) in the intracellular solution, and interleaved control experiments included DTT only.

A broken tip (∼10–20 μm) stimulating patch-type micropipette filled with ACSF was placed in the inner molecular layer (IML, < 50 μm from the border of the GC body layer) to activate MC axons, and in the middle third of the molecular layer to activate MPP inputs. For minimal stimulation (**Figure S6F and S6G**), stimulating pipettes were made from theta glass capillaries. To elicit synaptic responses, paired, monopolar square-wave voltage or current pulses (100–200 μs pulse width, 4-27 V) were delivered through a stimulus isolator (Digitimer DS2A-MKII). Typically, stimulation intensity was adjusted to obtain synaptic responses comparable in amplitude across experiments; e.g., 30-70 pA EPSCs (V_h_ −60 mV). For the optogenetic experiments, EPSCs were evoked using 1-3 ms pulses of blue (470-nm) light, provided by a collimated LED (Thorlabs, M470L3-C5, 470 nm, 300mW) and delivered through the microscope objective (40X, 0.8 NA). GC TBF-induced LTP was typically induced with 10 bursts at 5 Hz of 5 AP at 50 Hz, repeated 4 times every 5s, while the membrane potential was held at −60 mV in current-clamp mode. Extracellular field excitatory postsynaptic potentials (fEPSPs) were recorded using patch-type pipettes filled with 1M NaCl, and placed in *CA1 stratum radiatum*. fEPSPs were evoked with an ACSF-containing broken tip patch pipette in *CA1 stratum radiatum*. LTP at CA3-CA1 synapses was triggered using high frequency stimulation protocol (4 HFS: 100 pulses at 100 Hz repeated 4 times every 10 s) and recordings were performed at 25 ± 1°C. LTD was induced with low-frequency stimulation protocol (LFS: 900 pulses at 1 Hz).

Electrophysiological data were acquired at 5 kHz, filtered at 2.4 kHz, and analyzed using custom-made software for IgorPro (Wavemetrics Inc.). Paired-pulse ratio (PPR) was defined as the ratio of the amplitude of the second EPSC (baseline taken 1-2 ms before the stimulus artifact) to the amplitude of the first EPSC. Coefficient of variation (CV) was calculated as the standard deviation of EPSC amplitude divided by mean EPSC amplitude. Both PPR and CV were measured 10 min before and 20–30 min after LTP induction protocol or CGS21680-induced potentiation. PPR and CV were measured over minutes 0-6 before, 9-15 min after, and 29-35 min after CCPA application. The magnitude of LTP/LTD was determined by comparing 10 min baseline responses with responses 20-30 min (or 30-40 min for **Figure 1B**, 40-50 min for **Figure 2B**, 45-55 min for **Figure 2D**) after induction protocol. Averaged traces include 20 consecutive individual responses.

### Postsynaptic *TrkB* and *Bdnf* conditional KO

Adeno-associated virus AAV_5_.CamKII.eGFP (control virus) or AAV_5_.CamKII.GFP-CRE (University of Pennsylvania Vector Core) was injected (1 μL at a flow rate of 0.1 μL/min) unilaterally into the dorsal blade of the dentate gyrus (2.06 mm posterior to bregma, 1.5 mm lateral to bregma, 1.8 mm ventral from dura) of *TrkB_fl/fl_* or *Bdnf_fl/fl_* mice (4-5 week old). Animals were placed in a stereotaxic frame and anesthetized with isoflurane (up to 5% for induction and 1%–3% for maintenance). Slices for electrophysiology were prepared from injected animals 3–5 weeks after injection. For each animal, the absence of GFP-expressing cells in the hilus of the entire ipsilateral hippocampus was verified. We previously reported that this strategy can target GCs almost exclusively, as indicated by the lack of labeling in the hilus (Berthoux *et al*., 2023; Hashimotodani *et al*., 2017; Nasrallah *et al*., 2022). To conditionally KO *TrkB* from GCs, we also injected a control (ChiEFtom2A-GFP) or a Cre-expressing lentivirus selective for GCs (ChIEFtom2A-Cre, Post TrkB cKO) into the DG (relative to bregma: 1.9 mm posterior, 1.25 mm lateral, 2.3 ventral) of *TrkB_fl/fl_* mice (see **Figure S1C**). The lentivirus encoding ChiEFtom2A-Cre or ChiEFtom2A-GFP was under the control of the GC-specific C1ql2 promoter (Barthet *et al*., 2018), which was successfully used in our lab (Monday *et al*., 2022). In all cases, MC EPSCs were monitored in glowing GCs. C1ql2-ChiEFtom2A-GFP or C1ql2-ChiEFtom2A-Cre plasmids were a gift of Drs. Gaël Barthet and Christophe Mulle, Université de Bordeaux, and lentiviruses were made at the Einstein Genetic Engineering and Gene Therapy Core.

### Presynaptic *TrkB* and *Adora2a* conditional KO

A mix (1:2 ratio, 1.2 μL at 0.1 μL/min) of Cre recombinase-containing AAV (AAV_5_.CaMKII.Cre-mCherry, UNC Vector) and Cre-dependent ChiEF (AAV_DJ_.Flx.ChIEF.TdTomato) was injected into the dentate gyrus (relative to bregma: 1.9 mm posterior, 1.25 mm lateral, 2.3 ventral) of adult *TrkB_fl/fl_*, *Adora2a_fl/fl_* or WT control mice (4-5 week old). We then performed electrophysiology experiments 4–5 weeks post-injection in contralateral hippocampal slices. This allowed us to optically activate Cre-expressing MC axons.

### Optogenetic activation of supramammillary inputs

A beveled glass capillary pipette connected to a microsyringe pump (UMP3, WPI) was used for viral injection. 200 nL (50 nL/min) of AAV1-EF1a-DIO-hChR2(H134R)-eYFP (Addgene: 20298-AAV1) was injected into the SuM (relative to bregma, AP: −2.2 mm, ML: ±0.3 mm, DV: −4.85 mm) of VGluT2-Cre mice. The glass capillary remained at the target site for 5 min before the beginning of the injection and was removed 10 min after the infusion. Channelrhodopsin (ChR2)-expressing SuM axons were activated at 0.05 Hz by a pulse of 470 nm blue light (5 ms duration, 10.5 mW/mm_2_) delivered through a 40X objective attached to a microscope using an LED (Mightex).

### Pharmacology

Reagents were bath applied following dilution into ACSF from stock solutions stored at −20°C prepared in water, DMSO or ethanol, depending on the manufacturer’s recommendation. BDNF puffs (8 nM, 2.5-3 PSI, 3 s puffs repeated twice, 5 s interval) were applied using a Picospritzer III (Parker) connected to a broken patch pipette. The tip of the puffer pipette was positioned above the IML while monitoring MC-GC transmission. For experiments requiring postsynaptic loading PKI_6-22_, LTP was induced at least 20 min after establishing the whole-cell configuration. The time to the LTP induction was matched in interleaved controls.

### Electron Microscopy

Mice were anesthetized by intraperitoneal injection of ketamine⁄xylazine (ratio 1:1, 0.1 mL/kg) and transcardially perfused with an ice-cold fixative containing 4% paraformaldehyde, with 0.05% glutaraldehyde and 15% (v/v) saturated picric acid made up in 0.1 M phosphate buffer (PB, pH 7.4). Brains were then removed and immersed in the same fixative for 2 hours or overnight at 4°C. Tissue blocks were washed thoroughly in 0.1 M PB. Coronal sections (60-μm thick) were cut using a vibratome (Leica V1000). Pre-embedding immunohistochemical analyses were performed as described previously (Lujan et al., 1996). Free-floating sections were incubated in 10% (v/v) normal goat serum (NGS) diluted in Tris-buffered saline (TBS). Sections were then incubated in, 3-5 μg/mL diluted in TBS containing 1% (v/v) normal NGS, anti-A_2A_R [guinea pig anti-A_2A_R polyclonal (AB_2571656; Frontier Institute co., Japan)] or anti-A_1_R [rabbit-anti-A_1_R antibody (2 mg/ml; Affinity Bioreagents, Labome, USA)] antibodies, followed by incubation in goat anti-guinea pig IgG coupled to 1.4 nm gold or in goat anti-rabbit IgG coupled to 1.4 nm gold (Nanoprobes Inc., Stony Brook, NY, USA), respectively. Sections were postfixed in 1% (v/v) glutaraldehyde and washed in double-distilled water, followed by silver enhancement of the gold particles with an HQ Silver kit (Nanoprobes Inc.). Sections were then treated with osmium tetraoxide (1% in 0.1 m phosphate buffer), block-stained with uranyl acetate, dehydrated in a graded series of ethanol, and flat-embedded on glass slides in Durcupan (Fluka) resin. Regions of interest were cut at 70-90 nm on an ultramicrotome (Reichert Ultracut E, Leica, Austria) and collected on single-slot pioloform-coated copper grids. Staining was performed on drops of 1% aqueous uranyl acetate followed by Reynolds’s lead citrate. Ultrastructural analyses were performed in a Jeol-1010 electron microscope. Quantitative analysis of the relative abundance of A_2A_R or A_1_R, in the molecular layer of the dentate gyrus, was performed from 60 µm coronal slices as described (Lujan *et al*., 1996), in the area for MC-GC synapses and the area of PP-GC synapses. For each of the three animals, three samples of tissue were obtained (nine total blocks). Electron microscopic serial ultrathin sections were cut close to the surface of each block because immunoreactivity decreased with depth. Randomly selected areas were captured at a final magnification of 45,000X, and measurements covered a total section area of ∼5000 µm_2_. Dendritic shafts, dendritic spines and axon terminals were assessed for the presence of immunoparticles. The percentage of immunoparticles for A_2A_Rs or for A_1_Rs at postsynaptic and presynaptic sites was calculated. The perimeter of each axon terminal was measured in reference areas totaling ∼2,000 µm_2_. All axon terminals establishing excitatory synapses were counted and assessed from single ultrathin sections.

### Two-photon live imaging of GRAB_Ado_ in acute hippocampal slices

AAV_9_.hSyn.GRAB.Ado1.0m (WZ Biosciences Inc) was injected (1 μL at 0.1 μL/min) unilaterally into the hilus of WT mice or *TrkB^fl/fl^* mice (3 - 4 week old). Slices were prepared 3 - 4 weeks post-injection and expression of GRAB_Ado1.0m_ was confirmed in MCs of the ipsilateral hippocampus for each injected mouse. Hippocampal slices from GRAB_Ado1.0m_-injected mice were prepared using an ice-cold dissection buffer maintained in 5% CO_2_/95% O_2_ and containing (in mM): 25 NaHCO_3_, 1.25 NaH_2_PO_4_, 2.5 KCl, 0.5 CaCl_2_, 7 MgCl_2_, 25 D-glucose, 110 choline chloride, 11.6 ascorbic acid, 3.1 pyruvic acid. Slices were transferred to 32°C ACSF for 30 min and kept at room temperature for at least 45 min before imaging. Slices were then transferred to an imaging chamber under an Ultima 2-photon microscope (Bruker Corp.) equipped with a 60X NA 1.0 water-immersion objective and InSight DeepSee laser (Spectra-Physics). A 920-nm laser was used to excite GRAB_Ado1.0m_, and the emission signal was acquired using a 525–570 nm band-pass filter. The field of view (512 x 512 pixels per frame) was chosen in the IML of contralateral hippocampal slices where MC commissural axons projected onto GCs and expressed GRAB_Ado1.0m_. A broken tip stimulating patch-type micropipette filled with ACSF was positioned in the IML ∼150 μm from the imaging region to activate MC axons. The region of interest (ROI) was magnified to 2X and at least 40 consecutive images (at 0.25 Hz) as a baseline using PrairieView 5.4 (Bruker Corp.). Following baseline acquisition, MC burst stimulation was applied and at least an additional 100 images (at 0.25 Hz) were acquired. To verify the reactivity of the ROI, adenosine (100 μM) was added at the end of the imaging session. The fluorescence intensity of the ROI was measured and the ΔF/F_0_ of the GRAB_Ado1.0m_ signal was calculated using ImageJ software. All recordings were performed at 28 ± 1°C in a submersion-type recording chamber perfused at 2 mL/min with ACSF.

### Seizure induction and monitoring

Epileptic seizures were induced acutely in 2-3–month-old mice. Intraperitoneal (i.p.) injections of 20-30 mg/kg of kainic acid (KA, HelloBio HB0355) prepared in saline solution the same day were performed. For behavioral seizure scoring, mice were monitored during the 120 min post-injection, and behavioral seizures were scored, by an experimenter blind to condition (control *vs Adora2a* cKO), using a modified Racine scale as follows: stage 0: normal behavior, stage 1: immobility and rigidity, stage 2: head bobbing, stage 3: forelimb clonus and rearing, stage 4: continuous rearing and falling, stage 5: clonic-tonic seizure, stage 6: death. The maximum Racine score was recorded every 10 minutes and the cumulative seizure score was obtained by summing these scores across all 12 bins of the 120 min experiment.

### Fiber photometry in freely behaving mice

AAV_9_.hSyn.GRAB.Ado1.0m (1.2 μL at 0.1 μL/min) was unilaterally injected into the left DG (relative to bregma: 1.9 mm posterior, 1.25 mm lateral, 2.3 ventral) of *TrkB_fl/fl_* mice (5-6 week old). Contralaterally, AAV_5_.CamKII.eGFP (control) or AAV_5_.CamKII.GFP-CRE (*TrkB* cKO) was injected (0.5 μL at a flow rate of 0.1 μL/min) into the right DG (1.9 mm posterior, 1.25 mm lateral, 2.1 mm ventral). This allowed us to locally knockout *TrkB* from DG excitatory neurons. An optic fiber (200 μm diameter, NA = 0.37, Neurophotometrics) was then implanted unilaterally into the right DG above the IML (relative to bregma: 1.9 mm posterior, 1.25 mm lateral, 1.8 mm ventral) to record GRAB_Ado1.0m_ fluorescence *in vivo* where *TrkB* was KO as compared to control. 3-5 weeks after surgery, the fluorescence signal was monitored before and after KA i.p. injection (30 mg/kg). To record GRAB_Ado1.0m_ fluorescence, a three-channel multi-fiber photometry system (Neurophotometrics v1 Ltd) was used. 470 nm and 415 nm out-of-phase excitation lights were bandpass filtered and directed via a 20X objective (power: 30 μW). A single patch cord connected to the optic fiber implant was used to deliver light and collect the emitted fluorescence, which was filtered, and projected on a CMOS camera sensor. The open-source software Bonsai was used for data acquisition (40 frames/s rate). The fluorescence intensity profile of each channel was calculated as the mean pixel value of the region of interest. To calculate ΔF/F_0_, the 470 nm-evoked signal was normalized by the isosbestic signal (415 nm) to correct for photobleaching and potential artifacts. F_0_ corresponds to the average of the last three minutes (baseline) before KA injection. Optic fiber location was verified *post hoc*, to ensure that the fiber tip was in the ML of the DG. Animals with optic tips located outside of the DG or lacking AAV expression were excluded from the analysis. It is worth noting that the recording area is restricted to the vicinity (∼200 μm) under the fiber tip (Pisano et al., 2019).

### Post-hoc analysis of AAV expression and optic fiber location

Viral expression and optic fiber location were verified *post hoc*, at the end of the experiments (**Figure 7**). Mice were anesthetized with isoflurane (3-5%) and transcardially perfused with 4% paraformaldehyde (PFA) in 0.1 M sodium phosphate buffer (PBS). 50 µm-thick brain coronal sections were prepared using a DSK Microslicer (DTK-1000), stained with DAPI (1:1000) to label cell nuclei, and mounted with Prolong diamond antifade reagent montant (ThermoFisher) onto microscope slides. A Zeiss LSM 880 Airyscan Confocal microscope with Super-Resolution and ZEN (black edition) software and a 25X oil-immersion objective were used for the acquisition of all images in this study.

### Quantification and Statistical Analysis

The normality of distributions was assessed using the Shapiro-Wilk test. In normal distributions, Student’s unpaired and paired two-tailed t-tests were used to assess between-group and within-group differences, respectively. In unpaired t-tests, we checked for unequal variance. In cases where variance was unequal between normal distributions, we used a p-value that does not assume equal variance (Welch Correction). One-way ANOVA and One-way ANOVA repeated measure (RM) were used when more than two groups were compared. The non-parametric paired sample Wilcoxon signed rank test and Mann-Whitney’s U test were used in non-normal distributions. Statistical significance was set to p < 0.05 (∗∗∗ indicates p < 0.001, ∗∗ indicates p < 0.01, and ∗ indicates p < 0.05). All values are reported as the mean ± SEM. Statistical results are summarized in **Table S1**. All experiments included at least three animals per condition. Statistical analysis was performed using OriginPro software (OriginLab).

