## Supplementary Material for "Retrograde adenosine/A_2A_ receptor signaling facilitates excitatory synaptic transmission and seizures"

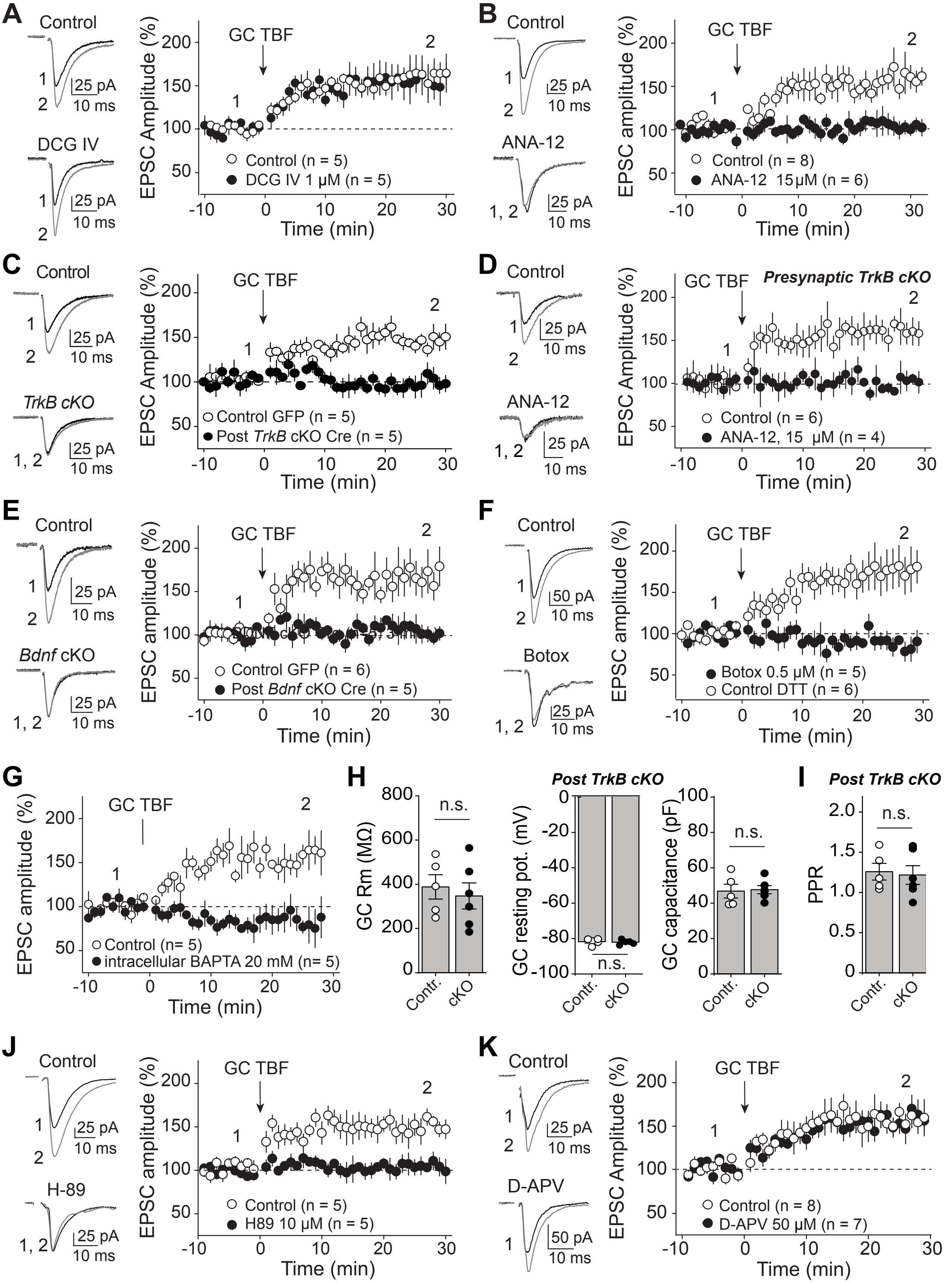

A
Control
DCG IV
25 pA
10 ms
GC TBF
EPSC Amplitude (%)
Time (min)
Control (n = 5)
DCG IV 1 μM (n = 5)
B
Control
ANA-12
GC TBF
EPSC amplitude (%)
Control (n = 8)
ANA-12 15 μM (n = 6)
C
Control
TrkB cKO
Control GFP (n = 5)
Post TrkB cKO Cre (n = 5)
D
Control
ANA-12
Presynaptic TrkB cKO
Control (n = 6)
ANA-12, 15 μM (n = 4)
E
Control
Bdnf cKO
Control GFP (n = 6)
Post Bdnf cKO Cre (n = 5)
F
Control
Botox
50 pA
Botox 0.5 μM (n = 5)
Control DTT (n = 6)
G
GC TBF
Control (n= 5)
intracellular BAPTA 20 mM (n= 5)
H
GC Rm (MΩ)
n.s.
Post TrkB cKO
GC resting pot. (mV)
GC capacitance (pF)
Contr.
cKO
I
Post TrkB cKO
PPR
J
Control
H-89
Control (n = 5)
H89 10 μM (n = 5)
K
Control
D-APV
Control (n = 8)
D-APV 50 μM (n = 7)

**Figure S1. Related to Figure 1: GC TBF-induced LTP properties**

**(A)** GC TBF induced a normal LTP in continuous presence of the group II mGluR agonist DCG-IV (1 μM) as compared to interleaved controls.

**(B)** TBF-induced LTP was blocked in continuous presence of the TrkB selective antagonist ANA-12 (15 μM).

**(C)** A control (ChiEFtom2A-GFP) or a Cre-expressing lentivirus selective for GCs (ChIEFtom2A-Cre, Post TrkB cKO) was injected into the DG of *TrkB*$^{fl/fl}$ mice. The lentivirus encoding ChiEFtom2A-Cre or ChiEFtom2A-GFP was under the control of the GC-specific C1ql2 promoter (Barthet et al., 2018). MC EPSCs were monitored in glowing GCs. LTP was abolished when TrkB was conditionally knocked out from GCs (Post TrkB cKO, *TrkB*$^{fl/fl}$ mice injected with ChiEFtom2A-Cre). LTP was unaffected in control animals (Control, *TrkB*$^{fl/fl}$ mice injected with ChiEFtom2A-GFP).

**(D)** Presynaptic TrkB cKO did not impair GC TBF-induced LTP. *TrkB*$^{fl/fl}$ mice were unilaterally injected with a mix of $AAV_5$.CaMKII.Cre and $AAV_{DJ}$.hSyn.Flex.ChiEF.Tdtomato. EPSCs were recorded in contralateral DG in response to light stimulation of Cre+ and ChIEF+ MC axons (lacking TrkB). GC TBF induced a normal LTP in presynaptic TrkB cKO mice, which was abolished in continuous presence of the TrkB antagonist ANA-12 (15 μM).

**(E)** GC TBF-induced LTP was blocked in GFP-Cre+ GCs (Bdnf cKO, *Bdnf*$^{fl/fl}$ mice injected with $AAV_5$.CaMKII.Cre.GFP) as compared with GFP+ GCs (control, *Bdnf*$^{fl/fl}$ mice injected with $AAV_5$.CaMKII.eGFP).

**(F)** Loading Botox (0.5 μM) in the postsynaptic neuron (GC) via the patch pipette completely abolished MC-GC LTP.

**(G)** Loading the calcium chelator BAPTA (20 mM) in the postsynaptic neuron (GC) via the patch pipette abolished GC TBF-induced LTP.

**(H)** No significant difference in GC membrane resistance (GC Rm), resting potential (GC resting pot.) or capacitance was found between control (*TrkB*$^{fl/fl}$ mice injected with $AAV_5$.CaMKII.eGFP) and postsynaptic TrkB cKO (*TrkB*$^{fl/fl}$ mice injected with $AAV_5$.CaMKII.Cre.GFP) mice.

**(I)** Basal MC-GC PPR was similar in control (*TrkB*$^{fl/fl}$ mice injected with $AAV_5$.CaMKII.eGFP) and TrkB cKO (*TrkB*$^{fl/fl}$ mice injected with $AAV_5$.CaMKII.Cre.GFP) animals.

**(J)** GC TBF failed to induce LTP in presence of the PKA inhibitor H89 (10 μM, 40- to 60 min pre-incubation and bath applied).

**(K)** Bath application of the selective NMDAR antagonist D-APV (50 μM) did not affect GC TBF-induced LTP, as compared with interleaved control experiments.

In A-F and J, K panels, representative traces are shown on the left and time-course summary plots on the right.
n.s. $p > 0.05$, unpaired t test.

Numbers in parentheses represent number of cells (n) and animals (N). Data are presented as mean ± SEM.

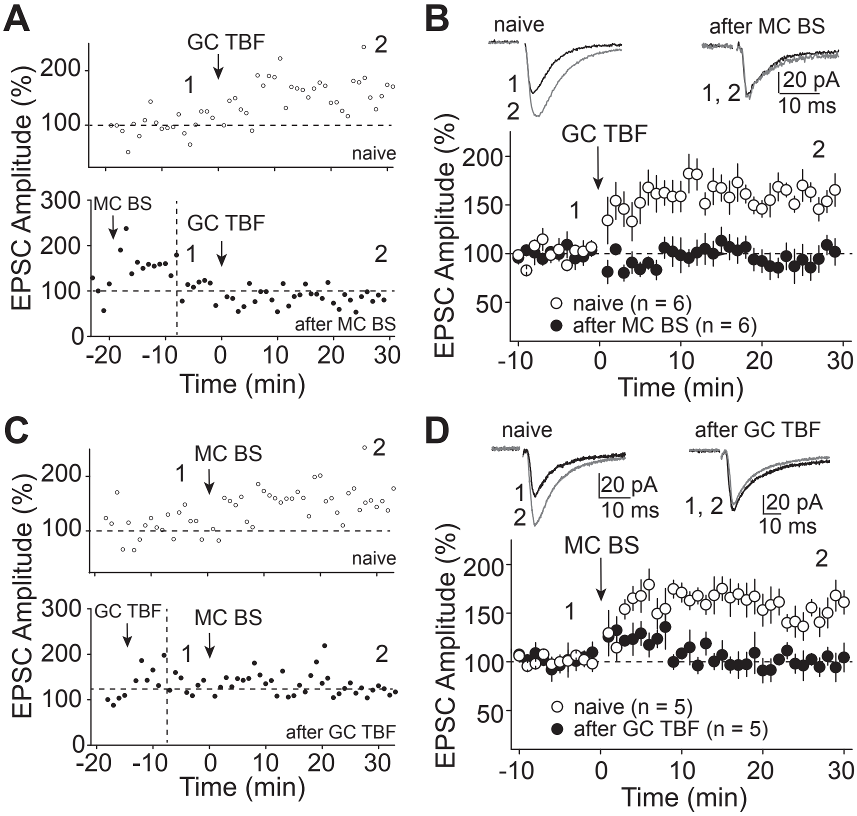


**Figure S2. Related to Figure 1: GC TBF- and MC burst stimulation (BS)-induced LTP occlude each other**

**(A)** Representative single cell experiments showing how GC TBF induced normal LTP in naive slices (top) but not when MC BS was previously applied (bottom). Vertical dashed line indicated the time point when stimulation intensity was reduced in order to avoid potential ceiling effect.

**(B)** Representative traces and time course summary plot showing how MC BS occluded GC TBF -induced LTP.

**(C)** Representative single cell experiments showing how MC BS induced a normal LTP in naive slices (top) but not when GC TBF was previously applied (bottom). Vertical dashed line indicated the time point when stimulation intensity was reduced in order to avoid potential ceiling effect.

**(D)** Pre-application of the GC TBF protocol occluded MC BS-induced LTP (black circles) as compared with interleaved controls (naive, white circles).

Numbers in parentheses represent number of cells. Data are presented as mean ± SEM.

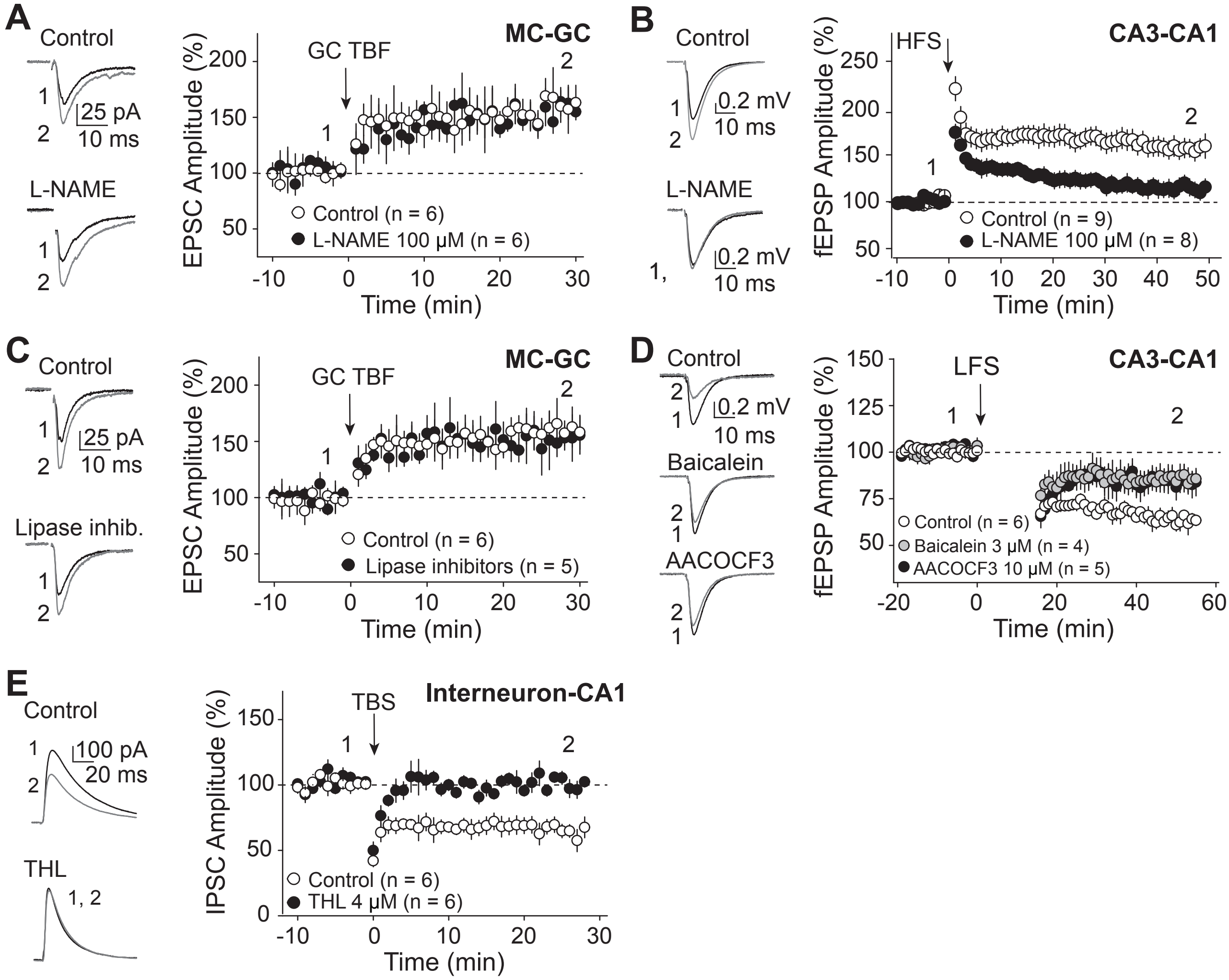


**Figure S3. Related to Figure 1: MC-GC LTP does not require conventional retrograde messengers**

**(A)** Bath application of the NO synthase inhibitor L-NAME (100 μM, 50- to 90-min pre-incubation and bath applied) did not impair MC-GC LTP.

**(B)** Positive controls for L-NAME. Extracellular recordings in CA1 stratum radiatum showing that L-NAME blocked LTP at CA3-CA1 synapses induced by 4 HFS (100 pulses at 100 Hz repeated 4 times every 10 s) at 25 °C.

**(C)** Application of cocktail of lipase inhibitors (10 μM of the fatty acid amide hydrolase (FAAH) and anandamide amidase inhibitor AACOCF3 and 3 μM of the lipoxygenases inhibitor Baicalein, 50- to 90-min pre-incubation and bath applied, and 4 μM of THL loaded in the recording pipette) did not affect MC-GC LTP.

**(D)** Positive controls for AACOCF3 and Baicalein. Extracellular recordings in CA1 stratum radiatum showing that application of AACOCF3 and Baicalein significantly reduced LFS (900 pulses at 1 Hz)-induced LTD at SC-CA1 synapses.

**(E)** Positive control for the lipase inhibitor THL showing that loading THL in CA1 pyramidal neuron via the recording pipette blocked iLTD induced by TBS (10 bursts at 5 Hz of 5 pulses at 100 Hz, repeated every 5 s, 4 times). Numbers in parentheses represent number of cells. Data are presented as mean ± SEM.

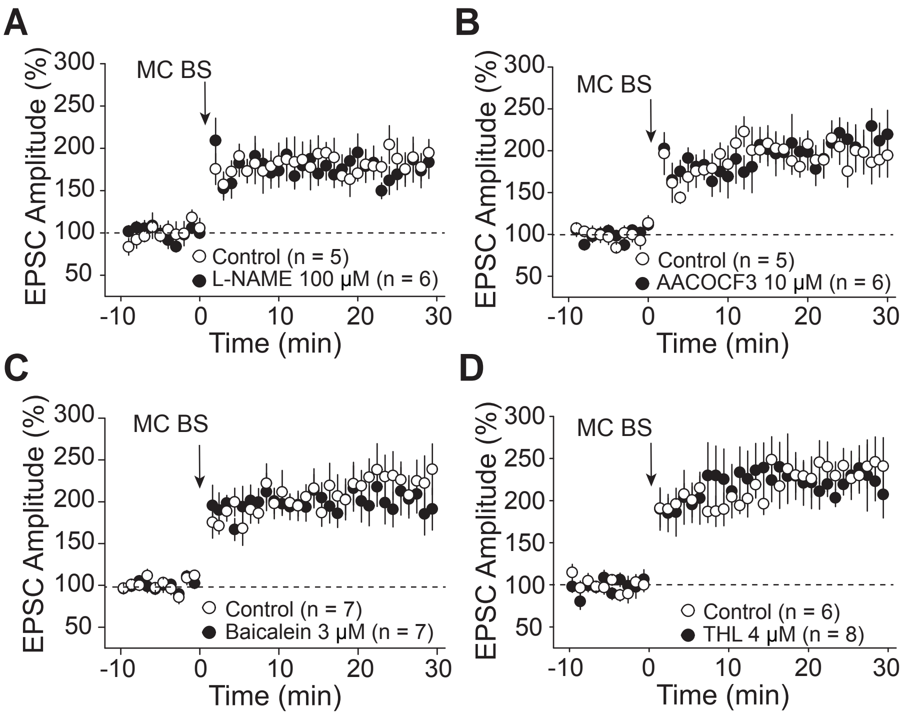


**Figure S4. Related to Figure 1: Synaptically-induced MC-GC LTP does not require conventional retrograde messengers**

MC BS induced a normal LTP in continuous presence of the NO synthase inhibitor L-NAME **(A)**, the fatty acid amide hydrolase (FAAH) and anandamide amidase inhibitor AACOCF3 **(B)**, the lipoxygenases inhibitor Baicalein **(C)**, and when the lipase inhibitor THL was loaded in the postsynaptic GC **(D)**, as compared with interleaved controls. Numbers in parentheses represent number of cells. Data are presented as mean ± SEM.

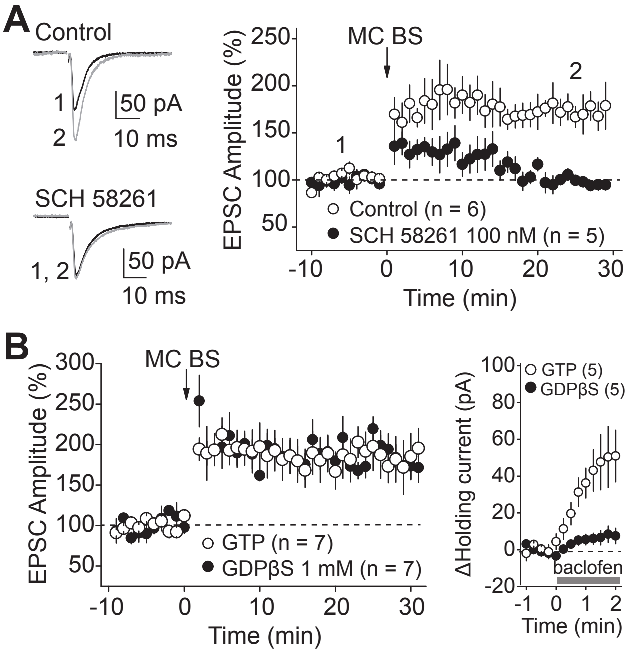


**Figure S5. Related to Figure 2: MC BS-induced LTP requires $A_{2A}$R activation**

**(A)** Bath application of the $A_{2A}$R selective antagonist SCH 58261 (100 nM) abolished LTP induced by MC BS (5 pulses at 100 Hz repeated 50 times every 0.5 s).

**(B)** Left, loading GDPβS (1 mM) in the recording pipette did not affect LTP induced by MC BS. Right, interleaved positive controls for GDPβS showing that replacing GTP by 1 mM GDPβS in the recording pipette abolished baclofen (10 µM)-induced increase in holding current. Baclofen is a $GABA_B$ receptor selective agonist.

Numbers in parentheses represent number of cells. Data are presented as mean ± SEM.

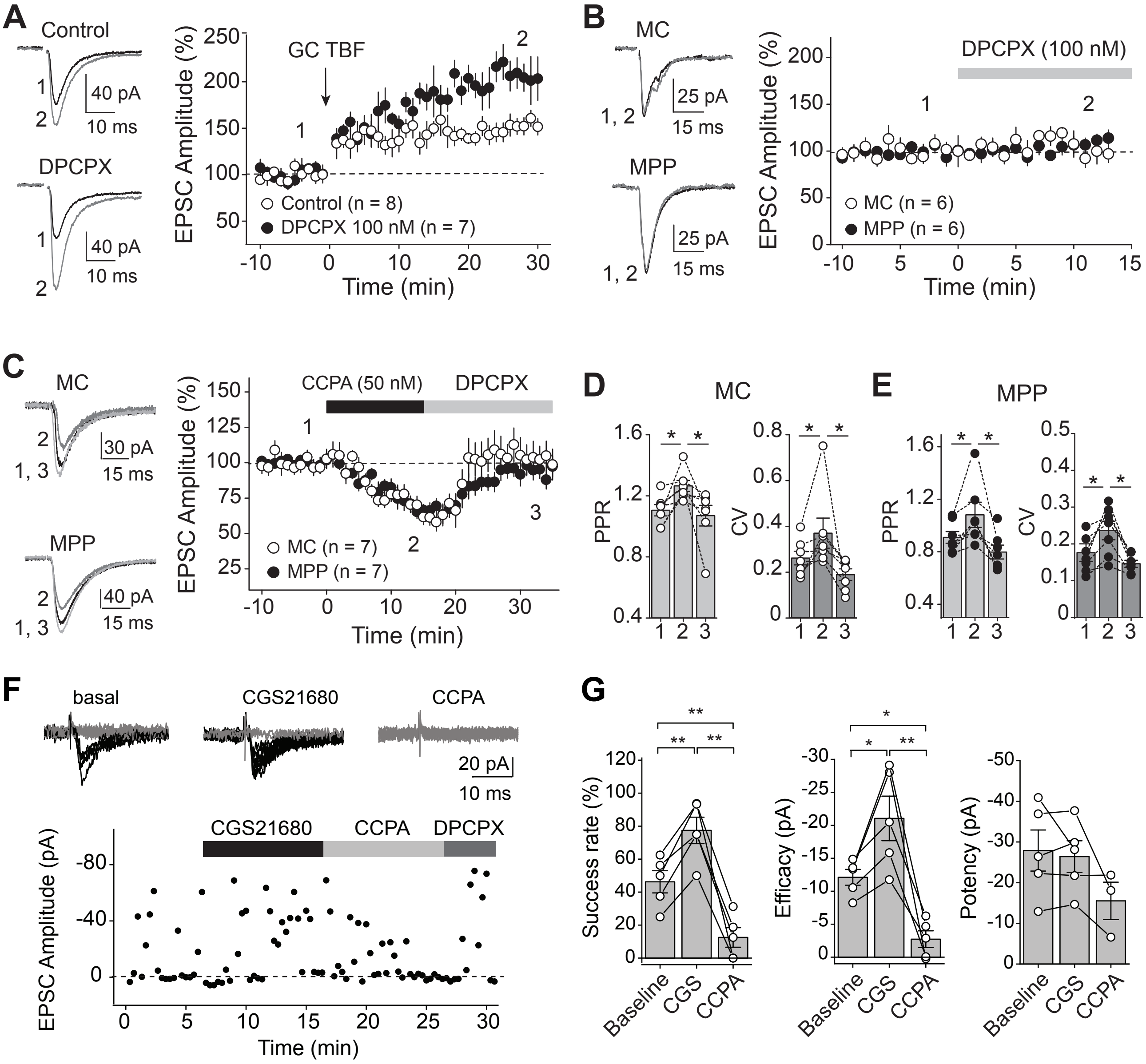


**Figure S6. Related to Figures 2 and 3: $A_1R$ activation dampens LTP but does not trigger LTD**

**(A)** MC-GC LTP magnitude was increased in the presence of the $A_1R$ selective antagonist DPCPX (100 nM) as compared to interleaved control experiments.

**(B)** Application of DPCPX (100 nM) did not change basal EPSC amplitude at MC-GC and MPP-GC synapses.

**(C)** Bath application of CCPA (50 nM, 15 min), a selective adenosine $A_1R$ agonist, induced a short-term depression at both MC-GC and MPP-GC synapses. The selective $A_1R$ antagonist DPCPX (100 nM) was included in the washout of CCPA.

**(D, E)** CCPA-induced depression was associated with a significant and reversible increase in both PPR and CV.

**(F)** Representative experiment using minimal stimulation in IML, time course (bottom) and sample traces (top).

**(G)** Summary plots demonstrating that $A_{2A}R$ agonist CGS21680 (50 nM) potentiated the synaptic responses evoked by minimal stimulation of MC axons, whereas subsequent application of the $A_1R$ agonist CCPA (50 nM) significantly reduced these potentiated responses recorded under the same conditions. Significant changes in success rate and efficacy are shown.

$*p < 0.05$, $** p < 0.01$. Numbers in parentheses represent number of cells. Data are presented as mean ± SEM.

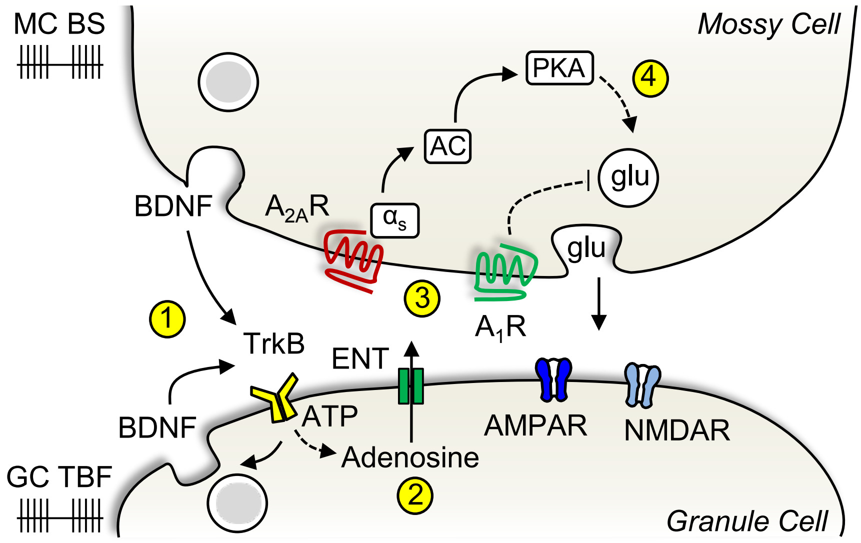


**Figure S7. Related to Figures 1-7: Adenosine is a retrograde messenger mediating activity-dependent presynaptic LTP at the MC-GC synapse**

Scheme illustrating the emerging model for the mechanism of activity-induced MC-GC LTP. MC (MC BS) and GC bursts (GC TBF) of activity trigger BDNF release and subsequent TrkB activation in GCs **(1).** Activation of postsynaptic TrkB leads to intracellular accumulation of adenosine, which is passively released from GCs via ENTs **(2)**. Adenosine then activates presynaptic $A_1$Rs and $A_{2A}$Rs **(3).** $A_{2A}$R activation induces a long-lasting PKA-dependent increase in glutamate (glu) release **(4)**, whereas $A_1$R activation dampens LTP.

**Table S1: Results and Statistical Tests Related to Figures 1-7 and S1-S6**

| Fig 1 | Results | P value | N | Test |
|---|---|---|---|---|
| Fig 1B | MC: 149.1 ± 7.1% of baseline<br>MPP: 100.0 ± 8.6% of baseline | MC: $p < 0.001$<br><br>MPP: $p > 0.9$ | MC: n = 7<br><br>MPP: n = 7 | MC: paired t-test<br><br>MPP: paired t-test |
| Fig 1C | PPR: n = 13, $p < 0.01$, paired t test;<br>CV: n = 13, $p < 0.01$, Wilcoxon signed rank test | PPR: $p < 0.01$<br><br>CV: $p < 0.01$ | PPR: n = 13<br><br>CV: n = 13 | PPR: paired t test<br>CV: Wilcoxon signed rank test |
| Fig 1D | *TrkB* cKO: 98.8% ± 2.6% of baseline<br>Control: 145.5% ± 7.7% of baseline | *TrkB* cKO: p = 0.8<br>Control: $p < 0.01$<br>*TrkB* cKO *vs* control: $p < 0.01$, | *TrkB* cKO: n = 5<br>Control: n= 5 | *TrkB* cKO: paired t-test<br><br>Control: paired t-test<br><br>*TrkB* cKO *vs* control: unpaired t test |
| Fig 1E | $PKI_{6\text{-}22}$ (pipette): 152.4% ± 10.2% of baseline<br><br>$PKI_{14–22}$ myristoylated (bath): 105.3% ± 4.1% of baseline | PKI6-22 (pipette): $p < 0.01$<br><br>PKI14–22 myristoylated (bath): p = 0.27 | PKI6-22 (pipette): n = 6<br><br>PKI14–22 myristoylated (bath): n = 5 | PKI6-22 (pipette): paired t test<br>PKI14–22 myristoylated (bath): paired t-test |
| Fig 1F | | ** $p < 0.01$<br><br>*** $p < 0.001$ | | paired t test |

| Fig 2 | Results | P value | N | Test |
|---|---|---|---|---|
| Fig 2A | control: 160.2% ± 8.9% of baseline,<br><br>SCH 58261: 97.4% ± 11.6% of baseline<br><br>ZM241385: 108.7% ± 3.7% of baseline | control: $p < 0.01$<br><br>SCH 58261: p = 0.83<br><br>control versus SCH 58261: $p < 0.001$<br><br>control versus ZM24138: $p < 0.05$ | control: n = 11<br><br>SCH 58261: n = 7<br><br>ZM241385: n = 5 | control: Wilcoxon signed rank;<br><br>SCH 58261: paired t test;<br><br>control versus SCH 58261: Mann-Whitney's U test<br>control versus ZM24138: Mann-Whitney's U test |
| Fig 2B | 102.9% ± 8.6% of baseline | p = 0.75 | n = 5 | paired t test |
| Fig 2C | control: 153.4% ± 7.4% of baseline<br>SCH 58261: 91.3% ± 5.9% of baseline | control: $p < 0.001$<br>SCH 58261: p = 0.22 | control: n = 7<br><br>SCH 58261: n = 5 | control: paired t test<br>SCH 58261: paired t test |

| | | | | |
|---|---|---|---|---|
| | | control versus SCH 58261: $p < 0.001$ | | control versus SCH 58261: unpaired t test |
| Fig 2D, left | control: 160.4% ± 10.6% of baseline,<br>GDPβs: 156.2% ± 10.2% of baseline | control: $p < 0.01$<br><br>GDPβs: $p < 0.01$, control versus GDPβs: $p = 0.78$ | control: n = 6<br><br>GDPβs: n = 6 | control: paired t test<br>GDPβs: $p < 0.01$ control versus GDPβs: unpaired t test |
| Fig 2D, right | control: 99.5 ± 18.3 pA<br><br>GDPβs: -12.5 ± 13.6 pA | control: $p < 0.05$<br><br>GDPβs: $p = 0.43$<br><br>control versus GDPβs: $p < 0.01$ | control: n = 4<br><br>GDPβs: n = 4 | control: paired t test<br>GDPβs: paired t test<br>control versus GDPβs: unpaired t test |
| Fig 2F | control: 151.9% ± 4.8% of baseline<br>Pre *Adora2a* cKO: 91.0% ± 18.9% of baseline | control: $p < 0.01$<br><br>Pre *Adora2a* cKO: $p = 0.29$<br><br>control versus Pre *Adora2a* cKO: $p < 0.01$ | control: n = 6<br><br>Pre *Adora2a* cKO: n= 6 | control: paired t test<br>Pre *Adora2a* cKO: paired t test<br>control versus Pre *Adora2a* cKO: unpaired t test |
| Fig 2G | control: 1.56 ± 0.7<br>Pre *Adora2a* cKO: 1.60 ± 0.8 | control versus Pre *Adora2a* cKO: $p = 0.67$ | control: n = 7<br><br>Pre *Adora2a* cKO: n= 6 | unpaired t test |

| **Fig 3** | **Results** | **P value** | **N** | **Test** |
|---|---|---|---|---|
| Fig 3A | MC: 152.1% ± 6.6% of baseline<br><br>MPP: 107.8% ± 5.2% of baseline | MC: $p <0.001$ 0.50<br><br>MPP: $p = 0.17$ | MC: n = 9<br><br>MPP: n = 9 | MC: paired t test<br><br>MPP: paired t test |
| Fig 3B | | PPR: $p < 0.01$<br><br>CV: $p < 0.01$ | PPR: n = 9<br><br>CV: n = 9 | PPR: paired t test<br>CV: Wilcoxon signed rank test |
| Fig 3C | MC: 153.1% ± 12.4% of baseline<br><br>SuM: 110.1% ± 11.0% of baseline | MC: $p < 0.01$<br><br>SuM: $p = 0.39$ | MC: n = 8<br><br>SuM: n = 9 | MC: paired t test<br><br>SuM: paired t test |
| Fig 3D | Pre *Adora2a* cKO: 91.1% ± 5.2% of baseline | Pre *Adora2a* cKO: $p = 0.16$ | Pre *Adora2a* cKO: n= 5 | Pre *Adora2a* cKO: paired t test |
| Fig 3E | control: 162.2% ± 6.4% of baseline<br>SCH 58261: 101.0% ± 5.3% of baseline | control: $p < 0.05$<br><br>SCH 58261: $p = 0.87$ | control: n = 5<br><br>SCH 58261: n = 5 | control: paired t test<br>SCH 58261: paired t test, |

| | | | | |
|---|---|---|---|---|
| | | control versus SCH 58261: $p < 0.05$ | | control versus SCH 58261: unpaired t test |
| Fig 3F | PKI$_{6-22}$ in the pipette: 160.5% ± 11.3% of baseline<br><br>PKI$_{14-22}$ in the bath: 98.6% ± 6.9% of baseline | PKI$_{6-22}$ in the pipette: $p < 0.01$,<br><br>PKI$_{14-22}$ in the bath: $p = 0.85$<br><br>PKI$_{6-22}$ in the pipette versus PKI$_{14-22}$ in the bath: $p < 0.01$ | PKI$_{6-22}$ in the pipette: n = 6,<br><br>PKI$_{14-22}$ in the bath: n = 5 | PKI$_{6-22}$ in the pipette: paired t test<br>PKI$_{14-22}$ in the bath: paired t test<br>PKI$_{6-22}$ in the pipette versus PKI$_{14-22}$ in the bath: unpaired t test |
| Fig 3G | Pre *Adora2a* cKO: 135.5% ± 4.7% of baseline | Pre *Adora2a* cKO: $p < 0.01$ | Pre *Adora2a* cKO: n= 5 | Pre *Adora2a* cKO: paired t test |
| Fig 3H | control: 153.6% ± 9.6% of baseline<br>ANA-12: 151.3% ± 4.5% of baseline | control: $p < 0.01$,<br><br>ANA-12: $p < 0.001$<br>control versus ANA-12: $p = 0.84$, | control: n = 6<br><br>ANA-12: n = 5 | control: paired t test,<br>ANA-12: paired t test,<br>control versus ANA-12: unpaired t test, |
| Fig 3I | control: 166.3% ± 8.3% of baseline<br>ANA-12: 101.2% ± 14.2% of baseline | control versus ANA-12: $p < 0.001$ | control: n = 6<br><br>ANA-12: n = 5 | control versus ANA-12: unpaired t test |
| Fig 3J | naive: 149.0% ± 6.3% of baseline<br>after CGS21680: 100.7% ± 9.2% of baseline | naive: $p < 0.001$<br><br>after CGS21680: $p = 0.44$<br>naive versus after CGS21680: $p < 0.001$ | naive: n = 6<br><br>after CGS21680: n = 4 | naive: paired t test,<br>after CGS21680: paired t test,<br>naive versus after CGS21680: unpaired t test, |
| Fig 3K | naive: 159.4% ± 4.6% of baseline<br>after CGS21680: 99.1% ± 11.6% of baseline | naive: $p < 0.001$<br><br>after CGS21680: $p = 0.89$<br>naive versus after CGS21680: $p < 0.001$ | naive: n = 5<br><br>after CGS21680: n = 4 | naive: paired t test,<br>after CGS21680: paired t test,<br>naive versus after CGS21680: unpaired t test, |

| **Fig 5** | **Results** | **P value** | **N** | **Test** |
|---|---|---|---|---|
| Fig 5A | control: 159.4% ± 9.9% of baseline<br>ENT inhibitors: 97.6% ± 8.7% of baseline | control: $p < 0.001$,<br><br>ENT inhibitors: $p = 0.80$,<br><br>control versus ENT inhibitors: | control: n = 7<br><br>ENT inhibitors: n = 5 | control: paired t test;<br>ENT inhibitors: paired t test; |

| | | | | |
|---|---|---|---|---|
| | | $p < 0.01$ | | control versus ENT inhibitors: unpaired t test |
| Fig 5B | 92.8% ± 7.2% of baseline | $p = 0.43$ | n = 6 | Wilcoxon signed rank test |
| Fig 5C | Control: 143.5% ± 6.3% of baseline<br>SCH 58261: 102.4% ± 4.1% of baseline | Control: $p < 0.01$<br><br>SCH 58261: $p = 0.59$<br>control versus SCH 58261: $p < 0.05$ | Control: n = 5<br><br>SCH 58261: n = 5 | Control: paired t test<br>SCH 58261: paired t test;<br>control versus SCH 58261: unpaired t test |
| Fig 5D | control: 158.4% ± 13.1% of baseline<br>inosine: 97.6% ± 4.1% of baseline | control: $p < 0.05$,<br><br>inosine: $p = 0.60$,<br><br>Control vs inosine: $p < 0.01$ | control: n = 5<br><br>inosine: n = 5 | control: paired t test,<br>inosine: paired t test,<br>control versus inosine: unpaired t test |
| Fig 5E | 96.3% ± 5.6% of baseline | $p = 0.55$, paired t test | n = 5 | paired t test |
| Fig 5F | Inhibitors in bath: 162.2% ± 19.2% of baseline<br><br>Inosine pipette: 97.6% ± 4.1% of baseline | Inhibitors in bath: $p < 0.05$<br><br>Inosine pipette: $p = 0.60$ | Inhibitors in bath: n = 5,<br><br>Inosine pipette: n = 5 | Inhibitors in bath: paired t test<br><br>Inosine pipette: paired t test |
| Fig 5G | control: 153.3% ± 5.4% of baseline<br>No ATP: 98.0% ± 1.9% of baseline | control: $p < 0.001$,<br><br>No ATP: $p = 0.36$<br><br>Control vs no ATP: $p < 0.0001$ | control: n = 6<br><br>No ATP: n = 4 | control: paired t test,<br>No ATP: paired t test,<br>control versus No ATP: unpaired t test |

| **Fig 6** | **Results** | **P value** | **N** | **Test** |
|---|---|---|---|---|
| Fig 6F | | Control vs ANA-12: $p < 0.05$<br>Control vs SCH: $p < 0.01$<br>Control vs TrkB cKO: $p < 0.001$<br>ANA-12 vs SCH: $p = 0.97$<br>ANA-12 vs cKO: $p = 0.58$<br>SCH vs cKO: $p = 0.86$<br>Control vs BDNF cKO: $p < 0.01$ | Control: n = 9<br>SCH: n = 7<br>ANA-12: n = 8<br>TrkB cKO: n = 8<br>BDNF cKO: n = 8 | One-way ANOVA |

| **Fig 7** | **Results** | **P value** | **N** | **Test** |
|---|---|---|---|---|

| | | | | |
|---|---|---|---|---|
| Fig 7A | control: 0.80 ± 0.08<br>Pre *Adora2a* cKO: 1.46 ± 0.16 | Control vs Pre *Adora2a* cKO: p < 0.01 | Control: n = 5<br>Pre *Adora2a* cKO: n = 4 | unpaired t test |
| Fig 7F, Latency to stage 3 (convulsion) | control: 24.5 ± 2.5 min<br>*Adora2a* cKO: 57.7 ± 10.6 min | Control vs *Adora2a* cKO: p < 0.01 | Control: n = 11<br>$A_{2A}$R KO: n = 13 | Mann-Whitney's U test |
| Fig 7F, Sum score | control: 35.4 ± 3.8<br>*Adora2a* cKO: 31.2 ± 4.8 | Control vs *Adora2a* cKO: p = 0.50 | Control: n = 11<br>$A_{2A}$R KO: n = 13 | unpaired t test |
| Fig 7H, Latency to stage 3 (convulsion) | control: 25.0 ± 5.0<br><br>*TrkB* cKO: 80.0 ± 16.7 | Control vs *TrkB* cKO: p < 0.05 | Control: n = 6<br>*TrkB*: n = 5 | unpaired t test |
| Fig 7H, sum score | control: 37.3 ± 6.5<br>*TrkB* cKO: 17.4 ± 3.6 | Control vs *TrkB* cKO: p < 0.05 | Control: n = 6<br>*TrkB*: n = 5 | Mann-Whitney's U test |
| Fig 7L | Control (post KA): 0.04 ± 0.01<br><br>*TrkB* cKO (post KA): 0.01 ± 0.003 | Control: p < 0.01<br><br>*TrkB* cKO: p = 0.06<br><br>Control vs *TrkB* cKO: p < 0.01 | Control: n = 4<br><br>*TrkB*: n = 5 | Control: paired t test<br><br>*TrkB* cKO: paired t test<br><br>Control vs *TrkB* cKO: unpaired t test |

| **Fig S2** | **Results** | **P value** | **N** | **Test** |
|---|---|---|---|---|
| Fig S2B | naive: 159.0% ± 7.3% of baseline<br>after MC BS: 91.5% ± 11.7% of baseline | naive: p < 0.001<br><br>after MC BS: p = 0.50<br>naive versus after MC BS: p < 0.001 | naive: n = 6<br><br>after MC BS: n = 6 | naive: paired t test,<br>after MC BS: paired t test,<br>naive versus after MC BS: unpaired t test, |
| Fig S2D | naive: 151.3% ± 9.5% of baseline<br>after GC TBF: 96.9% ± 8.2% of baseline | naive: p < 0.01<br><br>after GC TBF: p = 0.73<br><br>naive versus after GC TBF: p < 0.01 | naive: n = 5<br><br>after GC TBF: n = 5 | naïve: paired t test;<br>after GC TBF: paired t test;<br><br>naive versus after GC TBF: unpaired t test |

| **Fig S3** | **Results** | **P value** | **N** | **Test** |
|---|---|---|---|---|
| Fig S3A | control: 156.9% ± 13.8% of baseline<br><br>L-NAME: 152.4% ± 5.6% of baseline | control: p < 0.01,<br><br>L-NAME: p < 0.001, | control: n = 6<br><br>L-NAME: n = 5 | control: paired t test;<br><br>L-NAME: n = 5 |

| | | | | |
|---|---|---|---|---|
| | | control versus L-NAME: p = 0.78, | | |
| Fig S3B | control: 157.0% ± 9.9% of baseline<br><br>L-NAME: 114.1% ± 8.4% of baseline | control: p < 0.001<br><br>L-NAME: p = 0.14<br><br>control versus L-NAME: p < 0.01 | control: n = 9<br><br>L-NAME: n = 8, | control: paired t test,<br>L-NAME: paired t test,<br>control versus L-NAME: unpaired t test |
| Fig S3C | control: 158.9% ± 12.2% of baseline,<br>lipase inhibitors: 150.7% ± 16.4% of baseline | control: p < 0.01<br><br>lipase inhibitors: p < 0.05<br><br>versus inhibitors: p = 0.69 | control: n = 6<br><br>lipase inhibitors: n = 5 | control: paired t test;<br>lipase inhibitors: paired t test;<br><br>control versus inhibitors: unpaired t test |
| Fig S3D | control: 63.8% ± 3.4% of baseline<br><br>AACOCF3: 84.5% ± 6.5% of baseline<br><br>Baicalein: 86.0% ± 5.2% of baseline, n = 4, control | control: p < 0.001,<br><br>control versus AACOCF3: p < 0.05<br><br>Baicalein vs control: p < 0.05 | control: n = 6<br><br>AACOCF3: n = 5,<br><br>Baicalein: n = 4 | Control: paired t test,<br><br>AACOCF3: paired t test,<br><br>control versus AACOCF3: unpaired t test<br><br>Baicalein vs control: unpaired t test |
| Fig S3E | control: 64.0% ± 6.1% of baseline<br><br>THL: 96.7% ± 4.2% of baseline | control: p < 0.01,<br><br>THL: p = 0.46<br><br>control versus THL: p < 0.01 | control: n = 6<br><br>THL: n = 6 | control: paired t test<br><br>THL: paired t test<br><br>control versus THL: unpaired t test |

| **Fig S4** | **results** | **P value** | **N** | **Test** |
|---|---|---|---|---|
| Fig S4A | | control vs L-NAME: p > 0.05 | control: n = 5<br>L-NAME: n = 6 | unpaired t test |
| Fig S4B | | control vs AACOCF3: p > 0.05 | control: n = 5<br>AACOCF3: n = 6 | unpaired t test |
| Fig S4C | | control vs Baicalein: p > 0.05 | control: n = 7<br>Baicalein: n = 7 | unpaired t test |

| | | | | |
|---|---|---|---|---|
| Fig S4D | | control vs THL: $p > 0.05$ | control: n = 6<br>Baicalein: n = 8 | unpaired t test |

| **Fig S5** | **results** | **P value** | **N** | **Test** |
|---|---|---|---|---|
| Fig S5A | control: 174.5% ± 13.2% of baseline<br><br>SCH 58261: 97.9% ± 5.4% of baseline | control: $p < 0.01$,<br><br>SCH 58261: $p = 0.72$<br><br>control versus SCH 58261: $p < 0.001$ | control: n = 6<br><br>SCH 58261: n = 5 | control: paired t test<br><br>SCH 58261: paired t test<br><br>control versus SCH 58261: unpaired t test |
| Fig S5B, left | control: 185.8% ± 25.6% of baseline<br><br>GDPβs: 186.7% ± 29.4% of baseline | control: $p < 0.05$,<br><br>GDPβs: $p < 0.001$,<br><br>control versus GDPβs: $p = 0.35$ | control: n = 7,<br><br>GDPβs: n = 6 | control: Wilcoxon signed rank test;<br><br>GDPβs: paired t test;<br>control versus GDPβs: Mann-Whitney's U test |
| Fig S5B, right | | control versus GDPβs: $p < 0.05$ | control: n = 5<br>GDPβs: n = 5 | control versus GDPβs: unpaired t test |

| **Fig S6** | **results** | **P value** | **N** | **Test** |
|---|---|---|---|---|
| Fig S6A | control: 148.4% ± 6.4% of baseline,<br>DPCPX: 204.1% ± 15.9% of baseline | control versus DPCPX: $p < 0.05$ | control: n = 8<br><br>DPCPX: n = 7 | control versus DPCPX: unpaired t test |
| Fig S6B | MC: 105.2% ± 7.2% of baseline<br>MPP: 108.5% ± 7.7% of baseline | MC: $p = 0.50$<br><br>MPP: $p = 0.32$ | MC: n = 6<br><br>MPP: n = 6 | MC: paired t test;<br>MPP: paired t test |
| Fig S6C, MC | CCPA: 76.1% ± 4.6% of baseline<br><br>washout: 100.3% ± 26.8% of baseline | baseline versus CCPA: $p < 0.05$<br><br>washout versus CCPA: $p < 0.05$,<br><br>baseline versus washout: $p = 0.99$ | n = 7 | One-way ANOVA RM |
| Fig S6C, MPP | CCPA: 69.3% ± 4.6% of baseline<br><br>washout: 98.0% ± 7.6% of baseline | baseline versus CCPA: $p < 0.01$<br><br>washout versus CCPA: $p < 0.01$<br><br>baseline versus washout: $p = 0.96$ | n = 7 | One-way ANOVA RM |

| | | | | |
|---|---|---|---|---|
| Fig S6D,E | | * $p < 0.05$ | MC: n = 7<br>MPP: n = 7 | One-way ANOVA RM |
| Fig S6G, Success rate | Baseline: 46.2 ± 6.7<br>CGS21680: 77.4 ± 8.0<br>CCPA: 12.5 ± 5.9 | ** $p < 0.01$<br>*$p < 0.05$ | Baseline: n = 5<br>CGS21680: n = 5<br>CCPA: n = 5 | One-way ANOVA RM |
| Fig S6G, Efficacy | Baseline: -12.1 ± 1.2 pA<br>CGS21680: -21.1 ± 3.4 pA<br>CCPA: -2.7 ± 1.3 pA | ** $p < 0.01$ | Baseline: n = 5<br>CGS21680: n = 5<br>CCPA: n = 5 | One-way ANOVA RM |
